# Physical priors improve performance of structure-based binding affinity models

**DOI:** 10.64898/2026.09.11.750982

**Authors:** Benjamin Kaminow, Alexander Matthew Payne, Hugo MacDermott-Opeskin, John D. Chodera, Sukrit Singh

## Abstract

Structure-based drug discovery is a widely used paradigm for the rational design of novel small molecule therapeutics. However, the benefits conferred by the use of structural information has seen limited adoption in machine learning, where ligand-only ("2D") models are still the industry standard for molecular property or binding affinity prediction. Structure-based ("3D") ML models for binding-affinity prediction promise to present a clear advantage, but have not yet overtaken existing 2D models. Here, we show that physics-based priors can improve predictive performance of structure-based models by comparing different model architectures with varying physical priors on several prediction tasks. We present the Modular Training and Evaluation of Neural Networks (`mtenn`) package, where we decompose affinity prediction into separate steps of embedding structure into learned representations and combining those embeddings into a predicted binding affinity. We consider both E(3)-invariant and E(3)-equivariant architectures to determine the importance of encoding roto-translational inductive biases, as well as different methods for combining learned embeddings. By first optimizing several aspects of model construction using the general purpose PDBBind dataset, we are able to improve the performance and data efficiency of structure-based models. When subsequently trained and evaluated on the COVID Moonshot small molecule drug discovery dataset, our tuned models perform on par with industry standard ligand-only models. Our decomposed model framework highlights that encoding some physical priors improves model performance, while more complex biases such as equivariance offer limited benefit. Additionally, structure-based models generalize better to an unseen target and display higher training efficiency. Overall, these results emphasize that structure-based models benefit from their ability to incorporate physics-informed constraints, giving promising directions for model architecture development. These results also suggest that the strength of these models may be in tasks specifically aimed at generalizability, providing guidelines for their use in early-stage drug discovery campaigns.

## Introduction

Structure-based drug design (SBDD) is a widely used paradigm in small-molecule drug discovery, which has been accelerated by the introduction of computational tools [1–5]. SBDD operates on the premise that knowledge of a protein target’s structure should inform and accelerate the discovery of small molecule therapeutics [4, 6]. Several highly-utilized computational approaches that operate in this SBDD paradigm, including fragment merging, [7], docking or virtual screening to identify potent ligands [1, 8–10], molecular dynamics to simulate protein-ligand complex interactions [2], and free energy alchemical methods that calculate relative and absolute binding energies [11, 12]. Recently, there has been an explosive push of extending machine learning (ML) methods to use target-based structural information to inform the development of small molecule therapeutics [13–16]. A plethora of models have tackled the task of binding affinity prediction—predicting small molecule ligands’ equilibrium binding affinity to a target [13, 17–23]. However, many ML methods perform poorly in prospective contexts, often struggling with binding affinity prediction, generalizing to new targets, or posing compounds correctly [5, 24–28].

While one avenue to improve predictive utility is to increase the data available for training in the open research ecosystem through projects such as OpenBind [29], it remains valuable to identify promising architectures and model designs that better leverage existing datasets [5, 14, 30]. Ligand-only models have shown promising success in the task of binding affinity prediction [31–33]. Often referred to two-dimensional ("2D"), these models predict affinity using chemical structure/topology information of the candidate ligand only. Despite having access to less information than three-dimensional ("3D") models that predict affinity using structural data of the target and ligand, 2D models have often been found to perform at least as well as 3D models [27, 34]. The success of these 2D ML models is consistent with the long history of successful QSAR models and methods that have demonstrated useful predictive power in prospective discovery campaigns [31, 32]. However, there has been a persistent difficulty in demonstrating significant advantages to augmenting 2D ligand data with 3D target structural data [35, 36], which is counterintuitive given the long track record of successful use of structural information in physical models and other non-ML CADD tools.

Success in protein-ligand complex pose prediction by modern cofolding models demonstrates that structural data carries useful information, and suggests that model architectures can better use available structural information to improve predictive power [29, 37]. However, external benchmarks have shown that these cofolding models suffer from data leakage in their internal evaluations, potentially overstating their results and hindering widespread adoption and usage [24, 25, 28, 37–39]. Additionally, while some cofolding models are capable of predicting binding affinities alongside structure predictions, these predicted affinities have been found to be inaccurate [38, 40, 41]. Given these complications, the computational cost of training cofolding models presents a resource allocation question for the application of computational tools and resources towards design campaigns. In other words, training cheaper ligand-based models without the costly structural knowledge frees up computational resources to be allocated towards model deployment and drug design campaigns. Therefore, appropriate constraints and priors must be considered for structural models to become comparable to ligand-based models in accuracy and resource cost [5, 10, 42]. The inclusion of physical priors has been shown to be beneficial to ML model performance [15, 43]. Modern 2D ligand-based models have leveraged message passing graph neural networks (MPNNs) to learn representations from atomic identities and connectivities, rather than hand-encoding them in fingerprints [44–46]. MPNNs naturally encode atomic neighborhood information for each individual atom, and allow this information propagation to represent the interaction of functional groups in small molecules and amino acids. Three-dimensional models have shown great promise using representations like voxels, where structures are featurized into a 3D grid, or point cloud representations, where individual atoms are represented as nodes in a graph representation with edges based on interatomic distance [13, 35, 47, 48]. Point cloud representations in particular provide a generalizable encoding of atomic features and their relative coordinates [49]. Such representations have been found to be well suited to Graph Neural Networks (GNNs), and are well suited to modeling different levels of physical constraints such as E(3)-invariance and E(3)-equivariance. E(3)-invariant models are insensitive to the 3D Euclidean group of transformations (translations, rotations, and inversions), while E(3)-equivariant models are able to account for these operations [50–53]. Binding affinity is inherently E(3)-invariant in the protein-ligand complex coordinates; the relative translation, rotation, or inversion (provided an achiral solvent) of a protein-ligand complex in Euclidean space has no bearing on the binding affinity. As such, the inclusion of E(3)-invariance as an inductive bias aligns the model with the task of binding affinity prediction, theoretically reducing the scale of data needed or improving transferability of model predictive performance to new targets. Alternatively, using E(3)-equivariant representations within the model parallels the inter-atomic interaction forces these models will ideally learn to represent, which are naturally represented as vectors and therefore E(3)-equivariant themselves.

In this work, we assess how incorporating physics-based priors into structure-based ML models improves performance in binding affinity prediction relative to ligand-only models. We present the Modular Training and Evaluation of Neural Networks (`mtenn`) package, breaking structure-enabled affinity ML models into three modular components: *Representation*, *Strategy*, and *Readout*. This modular framework enables a reproducible fine-tuning of model construction and training practices, and facilitates assessing the impact that different choices have on model accuracy, data efficiency, and generalizability.

Evaluating against datasets like PDBBind [14] and the COVID Moonshot dataset [54], our modular strategy and subsequent inclusion of physics-based priors improves model performance. The formalization in `mtenn` also allows for careful hyperparameter tuning (Fig 1A), correcting for known instabilities in training 3D models and enabling them to perform on par with the baseline ligand-only models (Fig 4A). We find that pretraining of structure-based models on a general binding affinity dataset further improves their performance in the low-data regime (Fig 5B). Additionally, we evaluate the generalizability of 3D structure-based models by testing on antiviral data outside the COVID Moonshot or PDBBind, finding that models with structure-based priors perform better against unfamiliar targets. Lastly, we observe that these physical biases increase training efficiency, observing comparable performance in far fewer training epochs than 2D ligand-only models require. Through these analyses, we assess and identify where physical priors in structure-based ML models improve performance in binding affinity predictions.

**Figure 1.**
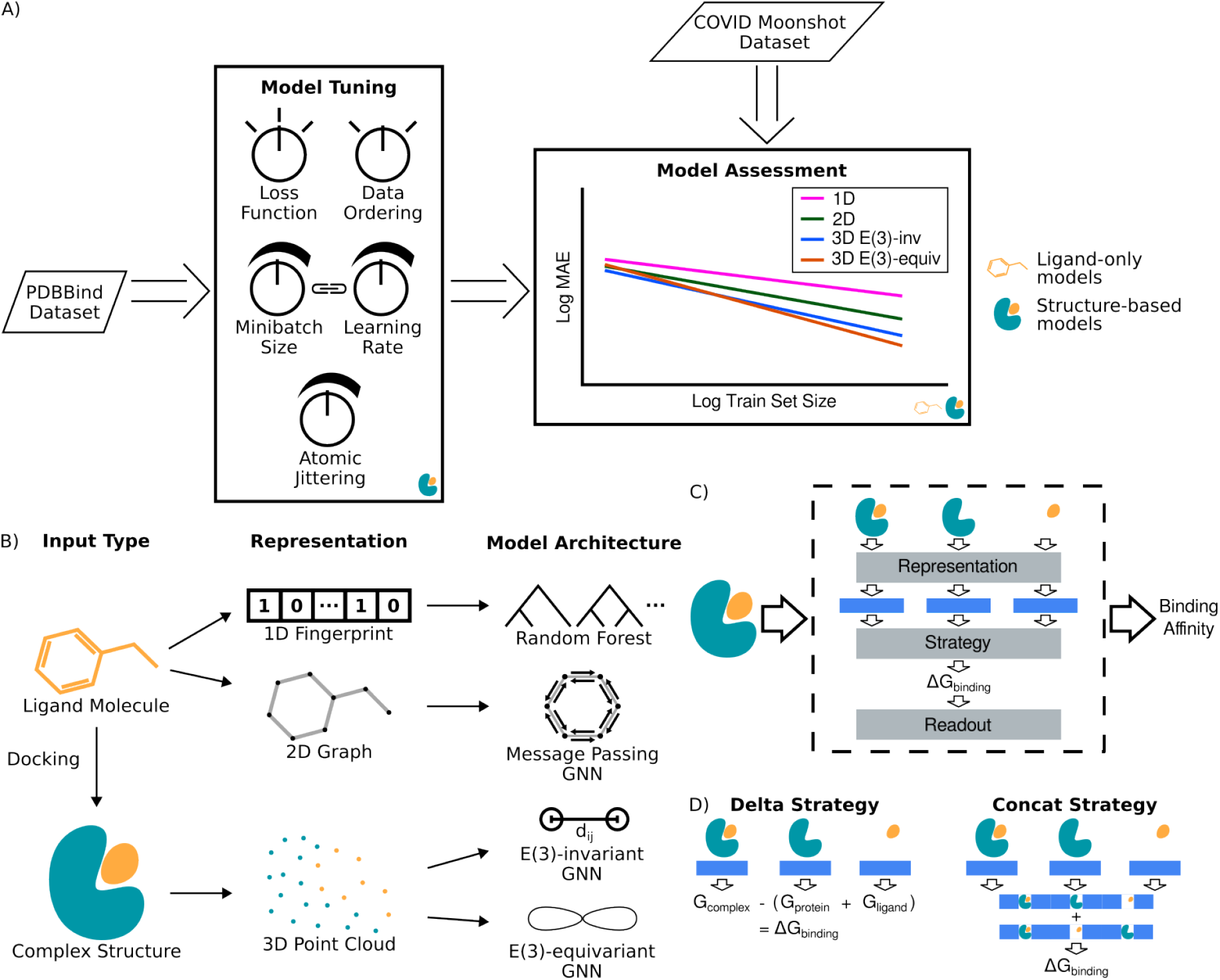
Using a modular approach to structure-based protein-ligand binding affinity predictions, we tune hyper-parameters using PDBBind before evaluating models on the COVID Moonshot dataset. A. Schematic representation of our workflow. We first perform hyperparameter optimization on structure-based models for binding affinity prediction using the general PDBBind dataset [14] (see **Hyperparameter Tuning on the PDBBind** in **Detailed Methods**). We then train and evaluate these tuned models on the COVID Moonshot [54] drug discovery dataset (Fig 2), comparing to industry standard ligand-only models as baselines. B. Machine learning representations for predicting binding affinity. Inputs to these models are either ligand-only information, such as molecular fingerprints (1D) or topology (2D), or proteinligand complex information, such as structure (3D) (described in Theory, **Methods, and Datasets**). C. Schematic of our decomposition of the affinity prediction task to optimize model construction and include physical priors. The proteinligand complex structure is first broken down to three substructures: the full complex structure, the protein structure alone, and the ligand structure alone. Each of these substructures is then independently passed as input to the same structure-based model (the *Representation* block), with the model predicting a latent space embedding for each of the substructures. These three predicted embeddings are passed together to the next layer (the *Strategy* block), which determines how these low-level representations are recombined to predict a binding free energy ΔG_binding_. Finally, this ΔG_binding_ is converted into an experimentally measurable binding affinity measurement readout, such as a pIC_50_ or pK_i_ (the *Readout* block). D. Two currently implemented algorithms for the *Strategy* block. The Delta *Strategy* uses a simple representation of the physical binding process, in which a simple linear neural network predicts an energy for each substructure and the resulting binding free energy ΔG_binding_ = G_complex_ - (G_protein_ + G_ligand_) is computed from bound and separated component free energies predicted by the model. The Concat *Strategy* allows the model to learn how these representations are combined by simply concatenating the learned representations, applying a sum-pooling layer, and allowing a linear neural network to predict a ΔG_binding_ directly from this concatenated embedding.

## Theory, Methods, and Datasets

### Different representations of input molecular data encode differing levels of information

Input data for ML models for binding affinity predictions can be represented using different levels of information, each encoding different priors into training (Fig 1B). More detailed representations, like structure, increase information provided for model training but also increase computational cost and the complexity of handling the data. [55] The least informative type of representation is the molecular fingerprint, in which a molecule is represented as a feature vector whose individual features describe chemical groups present in the molecule. [56, 57] Although efficient, these representations encode limited information about atom connectivity within the molecule. Atom graph representations address this shortcoming by representing atoms as nodes on a graph and bonds as edges connecting these nodes, ensuring that the molecule’s full connectivity is preserved. Molecular fingerprints and atom graphs are useful representations when there is no positional information to consider, however these formats are unable to encode structural data. In this case, three-dimensional representations are used, such as a voxel grid [13] or a 3D atomic graph [58, 59].

#### Ligand-only representations and models are competitive baselines for more information-rich data types

There are two main featurizations for computationally representing small molecules in ligand-only machine learning models: fingerprints and graphs (Fig 1B). Fingerprints are a one-dimensional (1D) representation that encode general information about the chemical makeup of a molecule, often represented as a bit vector encoding the presence or absence of various molecular features [56, 57, 60]. Models that operate on these 1D representations can be any traditional ML model that takes a feature vector as an input. In this work, we use LightGBM, a random forest-based architecture, as it is simple, easy to train, and interpretable [61, 62]. Additionally, previous efforts with LightGBM have demonstrated its suitability at handling sparser datasets and offering improved scalability with larger datasets [63, 64].

Graphs are a two-dimensional (2D) representation, which represent atoms in the molecule as nodes in a graph, and bonds as the edges between nodes (Fig 1B). These 2D graphs are also able to encode information about each atom and bond as node and edge features respectively [58]. Message passing graph neural networks are often used with 2D graphs, as they are able to take advantage of atom connectivity knowledge and allow atomic neighborhood information to propagate, which matches our understanding of chemistry and how functional groups interact within a molecule [45, 46, 65]. Through successive layers of updating node representations using the representations of neighboring nodes, each atom node representation contains information of not only the bonded neighbors, but the bonded neighbor of those nodes as well, with each layer extending the range of atoms included in a node’s representation. For this reason, it is important to balance the number of layers used in these models, as too few layers can lead to under-reaching, where a node’s information does not propagate beyond its immediate neighborhood, and too many layers can lead to over-smoothing, where the learned representations of all atoms are overly similar [66–68]. The number of layers must also be balanced with the size of the node feature embedding; as nodes compress information from larger numbers of neighbors into a fixed-length vector, it becomes more difficult for information from distant nodes to propagate (over-squashing).

Ultimately, we are generally interested in predicting a single value (e.g. binding affinity to a target protein) for the entire input graph, rather than returning the learned features for each node and edge. To calculate this final prediction, node features are pooled across all nodes to generate a single feature vector for the entire graph, which can then be fed to a standard feed-forward neural network [58]. In this work, we use a combination of weighted sum pooling and max pooling to calculate the graph-level feature vector as described in [69], in order to allow nodes to balance between learning from all neighbors and only the most important neighbor [48, 70, 71].

#### Structure-based representations and models contain more information but also more complexities

When expanding to 3D structures, the options for representation become more varied. One option is to generalize the 2D graph representation, adding coordinate information to each atom node in addition to its atom-related features [58, 59]. In the 3D graph case, edges are constructed between neighbors in physical space rather than between bonded atoms, and they encode a representation of this inter-atomic distance. These graphs can be constructed on an all-atom basis or at different levels of coarse-graining, for example representing each residue as a node rather than each heavy atom (Fig 1B). A non-graph-based approach to representing protein-ligand complex structures is using voxels, which discretizes space into a regular grid and assigns a value to each voxel based on the atomic content of that region of space. These voxel representations can then be used with standard convolutional neural network (CNN) approaches [13]. In this work, we exclusively use 3D point cloud representations, using all heavy atoms in both the protein and ligand and a one-hot encoding of each atom’s atomic number as the input node features. We exclude hydrogens from these representations as their positions fluctuate rapidly due to their weight, and are therefore difficult to resolve experimentally. Excluding hydrogens also reduces the computational cost and complexity for model training and handling with minimal information loss, as their positions can be inferred from heavy atom positions. Additionally, inclusion of hydrogens has consistently not been found to improve performance in empirical benchmarks [72].

Models operating on these 3D point cloud structure representations are often message passing neural networks, similar to the 2D case [58, 59, 73]. In these 3D representations, graph connectivity is built using physical position, treating all atoms within some cutoff as neighbors in the graph. In this work, we consider two different classes of 3D GNNs that represent different levels of physical priors: E(3)-invariant and E(3)-equivariant [43, 51, 52, 74]. E(3)-invariant models have internal representations that are invariant to the 3D Euclidean symmetry group (translations, rotations, and inversions), while E(3)-equivariant models are equivariant to the same [51, 52]. An E(3)-invariant model may be well-suited to predict binding affinities, as the target-ligand binding affinity is inherently E(3)-invariant; orientation of the protein-ligand complex has no bearing on the binding affinity. E(3)-equivariant representations may also be well suited to predicting binding affinities, as the underlying atomic interactions have inherent directionality. Although E(3)-equivariant models are more expressive, their usage of higher-order tensor math significantly increases the computational cost of training them. We build both classes of models using the `e3nn` package, using internal representations of *l* = 0 for the E(3)-invariant models and *l* = 1 for the E(3)-equivariant models (Fig 1B) [50, 51, 74–76]. By evaluating both E(3)-invariant and -equivariant models in this analysis, we evaluate whether the extra level of physical encoding has a positive effect on model performance, justifying the added computational cost.

### Decomposing structure-based prediction to enable modular evaluation of model construction

To facilitate optimizing and incorporating physical priors into 3D models, we decompose the task of structure-based binding affinity prediction into three steps (Fig 1C), which we formalize in the Modular Training and Evaluation of Neural Networks (`mtenn`) Python package. Within `mtenn`, this decomposition is organized into a *Model* class, which takes a protein-ligand complex structure as input, and returns a binding affinity prediction as an output. Within the *Model*, the input protein-ligand complex structure is first separated into its constituent substructures: the protein, the ligand, and the complex as a whole. This structural decomposition allows the model to learn how the different substructures interact; rather than attempting to predict a binding affinity based solely on a complex structure, the model can learn how to represent each individual substructure as well as how to combine these learned representations. After the structural decomposition, the *Model* contains three stages: the *Representation* block, *Strategy* block, and the *Readout* block, with the output of the *Readout* block being the final model prediction.

The first step of the model prediction is the *Representation* block, which predicts a low-level representation for each of the substructures (Fig 1C). This *Representation* block offers the first opportunity to adjust the model’s physical priors: by using underlying models that are E(3)-invariant or -equivariant, we can assess the importance of these inductive biases on binding affinity prediction. In `mtenn`, the *Representation* block is a thin wrapper around existing architectures. The minimal code requirement for adding a model as a *Representation* block allows for flexibility and efficiency when testing different architectures, as well as easy addition of novel model architectures.

The outputs from the *Representation* block are next fed to the *Strategy* block, which is responsible for combining these learned representation embeddings into a binding energy/affinity prediction(Fig 1C,D). The units of this prediction will depend on the final *Readout* block, as well as the units of the training data. In this work, we construct our models such that the output from our *Strategy* blocks are always ΔG_binding_ values in implicit kT units. This step of combining the learned representations also provides an opportunity for introducing physical biases in the system, and we evaluate two different *Strategy* implementations to assess the degree to which these physical biases are important at this step (Fig 1D). The Delta *Strategy* represents the thermodynamic cycle of ligand binding by predicting an energy for each of the three substructures and performing the definitional calculation of ΔG_binding_ = G_complex_ − (G_protein_ + G_ligand_) [12, 16]. The Concat *Strategy* is less constrained by prior knowledge of the equations governing binding, simply concatenating the three learned substructure embeddings and predicting the ΔG_binding_ directly from this combined feature vector. As a baseline, we use a complex-only strategy, that attempts to predict a ΔG_binding_ purely from the *Representation* output of the full complex structure. As with the *Representation* block, implementing new *Strategy* approaches is simple, and only requires wrapping the new method in a minimal code structure in order for it to be easily swapped into an existing `mtenn` model.

The final step in our decomposed model is the *Readout* step, which handles unit conversion of the output from the *Strategy* block into the units of the training data. Although this step is not strictly necessary, it ensures that the units of the *Strategy* block output are consistent. Currently, the *Readout* blocks implemented in `mtenn` expect a ΔG input in implicit kT units, which is then converted into a pK_i_ or pIC_50_ output. Importantly, the *Readout* block is not learned, which facilitates pretraining and the use of disparate data sets. By enforcing that the *Readout* block does not contain any model parameters, a model can be fully trained end-to-end on a dataset containing pK_i_ values, and subsequently used to predict pIC_50_ values simply by swapping the *Readout* block used. These *Readout* implementations make use of the Cheng-Prusoff equation [77]:

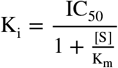

in conjunction with the definition of Gibbs free energy:

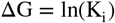

for ΔG in kT units, where [S] is the experimental substrate concentration and K_m_ is the Michaelis constant for this protein and ligand.

In this work, we use a combination of `mtenn`, which serves as the ML backend containing the models used, and the `drugforge-ml` package as our frontend, which contains machinery for reproducibly running and tracking model training. The modularity of these packages allows us to efficiently optimize hyperparameters of model construction and assess different methods of incorporating physical biases into the structure-based models.

We additionally use the rest of the `drugforge` package for structure preparation and ligand docking. Both `mtenn` and `drugforge` are installable via `conda-forge`, and their code is available on GitHub at https://github.com/choderalab/mtenn/ and https://github.com/choderalab/drugforge/.

### Evaluation against general purpose and specialized binding affinity datasets

#### The PDBBind is a large, general purpose dataset of protein-ligand complex structures and binding affinities

The PDBBind dataset contains a subset of the full Protein Data Bank that has been filtered down to contain only protein-ligand complex structures that were published alongside an experimentally measured binding affinity [14, 78]. The PDBBind is further divided into three subsets, based on structure resolution: the general subset, which contains the lowest-quality structures; the refined set, which contains structures that were collected based on a number of quality metrics; and the core set, which contains structures that were sampled from the refined set with the goal of reducing sample redundancy in the refined set. Additionally, the PDBBind core set serves as the primary test set of the CASF challenge [78]. The binding affinity measurements in the PDBBind are a mix of IC_50_, K_i_, and K_d_ values. Although these types of binding affinity measurements represent different biochemical properties [77, 79], Landrom and Riniker [80] show that IC_50_, K_i_, and K_d_ measurements available in the literature all show similar amounts of noise. We therefore combine these measurements, allowing us to construct the largest available dataset.

We use a leak-proof splitting of the PDBBind in order to minimize data leakage and appropriately assess model performance [81]. The PDBBind is commonly used as a dataset for biological machine learning, however its composition often leads to data leakage if the train, validation, and test sets are not constructed carefully [14]. Often, structures for the same protein are deposited to the PDB at different time points, meaning that a random or temporal split of the dataset would likely result in the same or similar proteins having structures in multiple different splits [26, 81, 82]. This data leakage can overstate model performance, leading to incorrect conclusions [83]. By using the leak-proof splitting described by Li, et al. [81], we ensure that the dataset splits that we use isolate structures from complexes in the same family or with similar function, with similar protein sequences, or with similar chemical structures. These leak-proof splits more accurately assess the models’ ability to extrapolate rather than interpolate, strengthening the decisions that we make based on these results.

#### The COVID Moonshot dataset provides a perfect test bed for structure-based ML in small-molecule drug discovery

In this study, we use the COVID Moonshot dataset to evaluate the models’ performance on a real drug discovery dataset [54]. Although the PDBBind is a useful dataset, its composition is not representative of a standard drug discovery campaign, meaning that a model that performs well on the PDBBind may not perform as expected in the context of a real drug discovery campaign [14, 81]. In contrast, the COVID Moonshot dataset is a large, high-quality, open drug discovery dataset generated during a real campaign targeting the SARS-CoV-2 main protease (Mpro) (Fig 2A) [54]. This dataset contains experimentally determined crystal structures of the SARS-CoV-2 Mpro in complex with several ligands and ligand fragments, as well as biochemical IC_50_ binding affinity measurements for over 1,000 different ligands and synthetic intermediates. Unlike in the PDBBind, all IC_50_ values in the COVID Moonshot dataset were generated consistently, using the same assay and with the same conditions, making them a reliable label to use in training a model.

**Figure 2.**
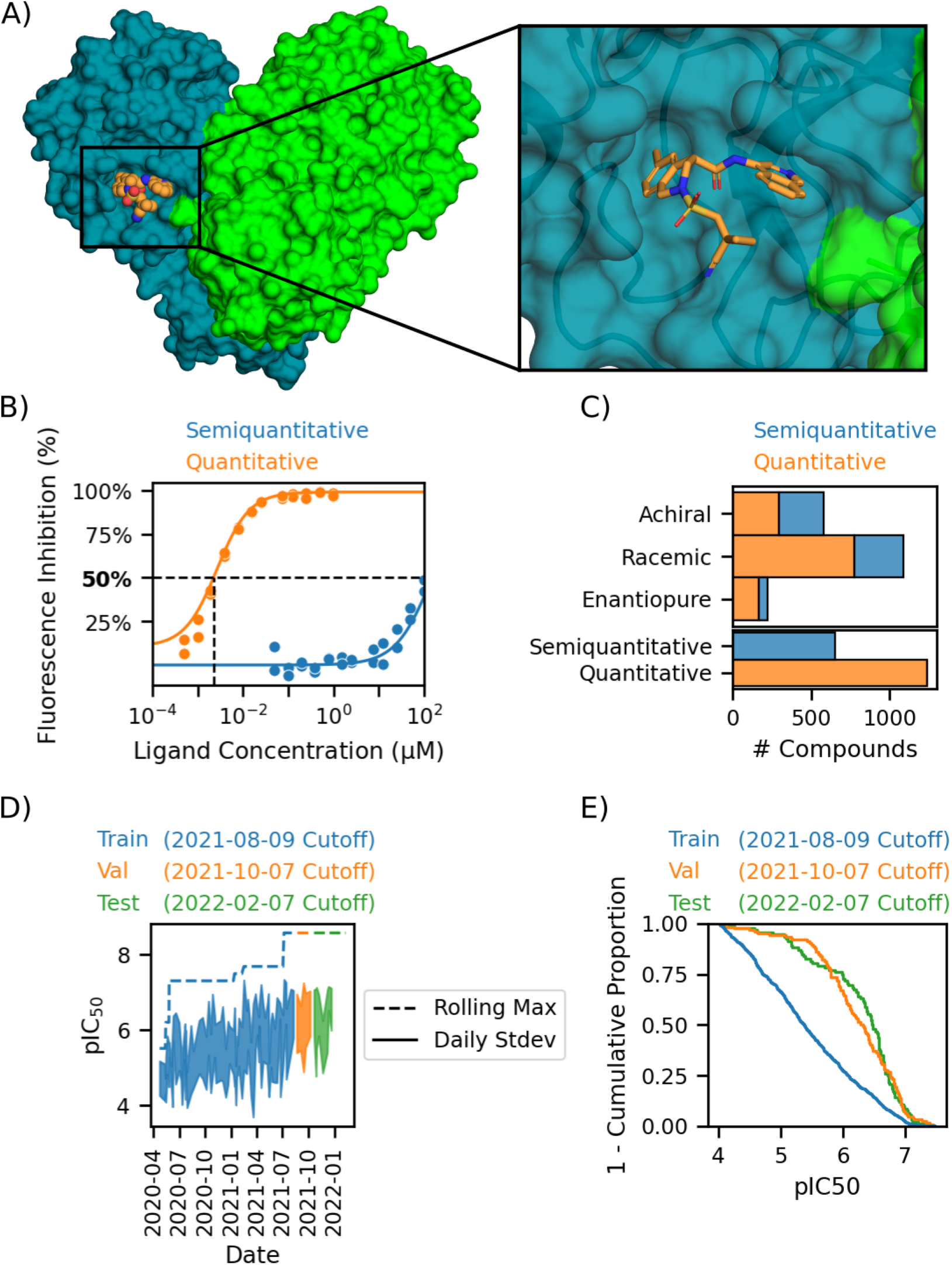
The COVID Moonshot curated dataset is a high-quality, time-stamped, open dataset from a real drug discovery program that can benchmark ML models in a real drug discovery context. A. Structure of SARS-CoV-2 main viral protease (Mpro; left, surface in turquoise and green; PDB ID 7GAW), the target for the COVID Moonshot open science discovery project. Shown here bound to COVID Moonshot molecule MAT-POS-e194df51-1 (orange spheres/sticks) [54]. A zoom insite of the binding site is shown (right). B. Example binding affinity curves of a molecule with a resolved (quantitative) IC_50_ (MAT-POS-7ed7af85-1, orange) and a molecule that is too weak of a binder to be resolved (semiquantitative, JAN-GHE-83b26c96-20, blue). These curves were generated using a fluorescence-based dose-response assay, with a significant fraction of the measurements lying outside of the assay dynamic range. C. Distribution of chemical matter types and measurement types in the COVID Moonshot dataset. Approximately half of the compounds are racemates, with 2/3 of the measurements being quantitative (orange). D. Time-based shift of pIC50 values over the course of the Moonshot campaign. As is typical in real-world drug discovery programs, compound potency (pIC_50_) undergoes a considerable distributional shift with time as more potent compounds are discovered. Both a rolling maximum (dashed line) and daily standard deviation (solid line and shaded region) are shown. E. Visualization of the temporal-splitting in our work. Due to the distributional shift that occurs as a drug discovery program identifies more potent molecules (D), we split our data temporally, taking earlier (likely to be less potent) molecules as the training and validation sets and the most recent (likely to be more potent) molecules as the test set for a realistic assessment of real-world performance in drug discovery programs.

The assay used to generate these binding affinity measurements was a fluorescence dose response assay (Fig 2B,C). Some ligands fall below the assay detection limit and are too weak of a binder for their affinity to be resolved, and some ligands exceed the assay detection limit and are too strong of a binder for their affinity to be resolved, making their measurements semiquantitative (e.g. > 100 *μ*M or < 1 nM). In both of these cases, although the measurement is less informative than one that falls within the assay limits, the measurement still encodes useful information. In our analyses, we adjust the loss functions and metrics used to account for this semiquantitative nature, counting a model prediction as correct if both the prediction and experimental value fall outside the assay limit in the same direction, i.e., either both values are above the assay limit or both are below the assay limit.

As is typical of a drug discovery campaign, the binding affinities trend towards stronger binders as the campaign progresses (Fig 2D) [5, 10, 84, 85]. To capture this trend in our model evaluation and ensure that model performance on this dataset is indicative of how these models would be used in a real campaign, we employ a temporal splitting to generate our train, validation, and test sets (Fig 2E) [10, 86]. For all experiments using the COVID Moonshot dataset, we use a fixed test set composed of the most recent 10% of compounds. To assess model data efficiency as well as overall performance, we train models on increasing amounts of data, always titrating compounds in from the most recent data. We always select the validation set as the 10% of compounds immediately following the train set.

## Results and Discussion

### Model hyperparameter tuning on the PDBBind improves model performance on the COVID Moonshot dataset

As our first step of model optimization, we perform hyperparameter tuning on our 3D models using a leak-proof splitting of the v2020 version of the PDBBind [14, 81]. This hyperparameter tuning consists of modifying 5 different aspects of model construction and training: the loss function, minibatch size, and learning rate used in training, whether the ordering of the training set was kept consistent each training epoch, and the application of positional jittering to the input training structures (Fig 3A). The full details of this hyperparameter tuning, including the relevant statistical analyses, are available in Detailed Methods. After this initial hyperparameter tuning, we assess the importance of introducing physical biases into the model by comparing the two *Strategy* methods (Fig 3B), as well as a baseline of using only the complex representation for the final binding affinity prediction. For this experiment, we evaluate model performance across all dataset sizes, as we are interested in both absolute model performance and model data efficiency. The complex-only baseline *Strategy* performs worse than either the Delta or Concat *Strategy*, which confirms the hypothesis that including physical priors by allowing the model to learn how the different constituent substructures interact is beneficial to model performance. We find that, overall, the more physically constrained Delta *Strategy* outperforms the Concat *Strategy* for both model architectures, and that all model/*Strategy* combinations display similar data efficiency dynamics (Fig 3B). In the low data regime, this trend is inverted for the E(3)-invariant model, however the Delta *Strategy* again outperforms the Concat *Strategy* after ∼500 training compounds.

**Figure 3.**
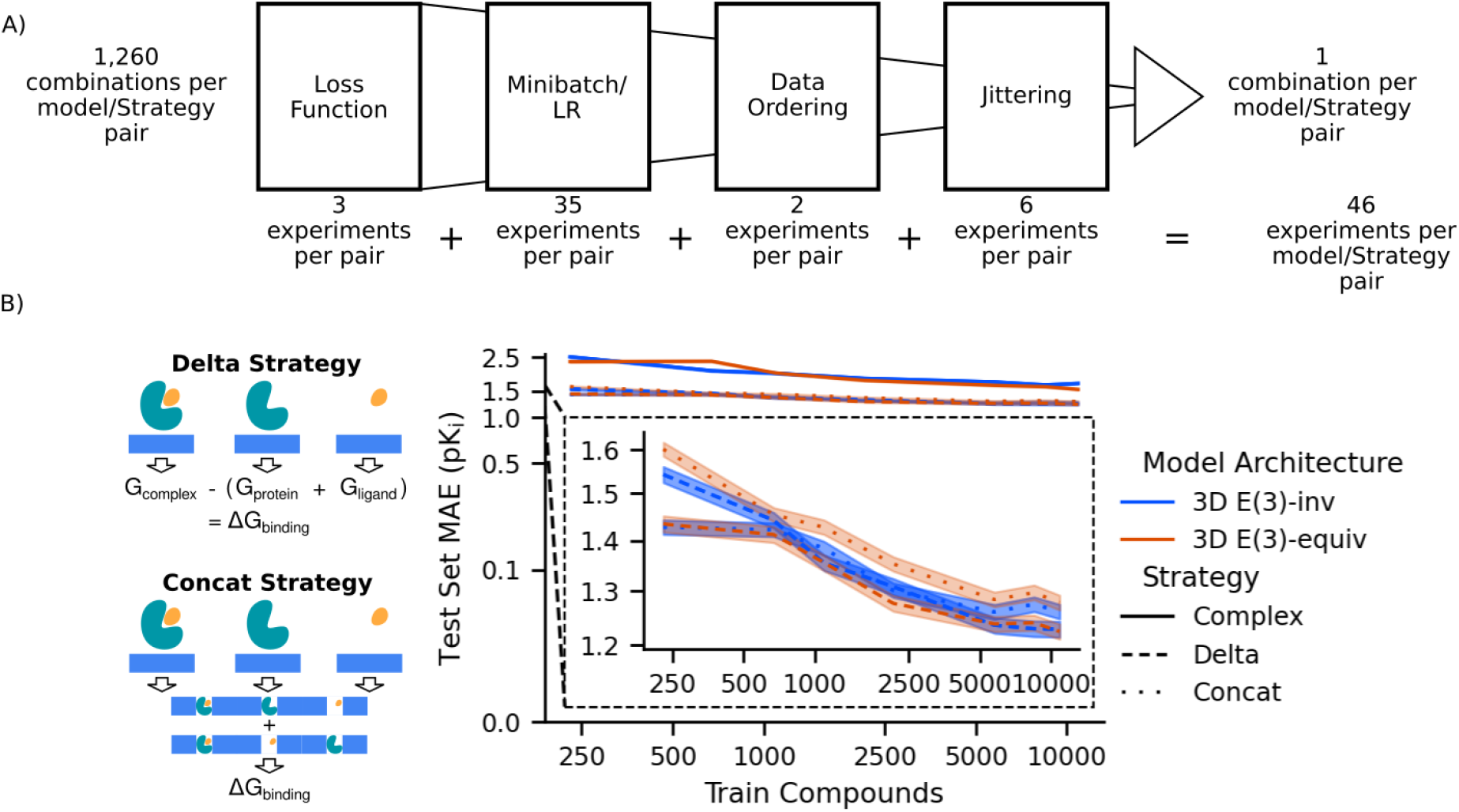
A sequential approach to optimizing model construction focuses computational scope while uncovering insights into structure-based modeling for binding affinity prediction. A. Schematic of sequential hyperparameter tuning. For each combination of *Strategy* and 3D model architecture, we evaluate and fix the choice of loss function used in training (left), the combination of learning rate and minibatch size (second from left), the order in which training data is shown to the model each epoch (second from right), and the application of atomic positional jittering as a regularization approach (right). We select the highest performing option for each of these hyperparameters for each model/*Strategy* pair sequentially, fixing each one when evaluating subsequent hyperparameters. The number of experiments per pair is shown below each corresponding step. The full table of evaluated hyperparameters is shown in **Table S1**, and results for these tuning experiments are shown in **Figs S1-S4.** B. Comparison of the two *Strategy* choices. Both the Delta *Strategy* (dashed line) and Concat *Strategy* (dotted line) significantly outperform a complex-only *Strategy* (solid line). When comparing between the *Strategy* choices that incorporate the representations from all three substructures, the Delta *Strategy* outperforms the Concat *Strategy* in terms of data efficiency and model performance across both models.

This improvement with the Delta *Strategy* relative to the Concat *Strategy* is especially notable in the E(3)-equivariant model, implying that these types of models may be able to extract more information from the physical intuition than E(3)-invariant models. However, the E(3)-invariant model with the Delta *Strategy* still performs on par with the E(3)-equivariant model with the Delta *Strategy*, so it is also possible that this difference in performance improvement comes simply from the E(3)-invariant model being closer to reaching a theoretical performance-maximum given the training dataset. Although the improvement varies in effect size, it is shared across both architectures, so we use the Delta *Strategy* in all 3D models in future experiments.

We evaluate the hyperparameter-tuned models on the COVID Moonshot dataset, and compare the models’ performance to that of their default parameter equivalents (Fig 4A). We use only the achiral and enantiopure molecules from the COVID Moonshot dataset, as it is not straightforward to handle racemic data with structure-based models. The Fragalysis crystal structure repository is publicly accessible, and the COVID Moonshot molecules and associated relevant data are available in the data repository that accompanies this manuscript (see Code and Data Availability). We generated the structures used for dataset generation by docking each molecule to the crystal structure whose ligand shares a maximal common substructure with the ligand to be docked, selecting the top-ranked pose for each protein-ligand complex. Payne, et al. have shown that when there is high enough similarity between candidate ligands and available cocrystalized ligands as references, docking is able to successfully capture true binding modes [10]. Additionally, this work uses the same dataset used in [10], so given the success of docking shown therein we believe our docked structures serve as a reasonable starting point. This MCS search and the subsequent docking using the OpenEye POSIT algorithm was performed using the `drugforge` software package [1, 42, 87].

**Figure 4.**
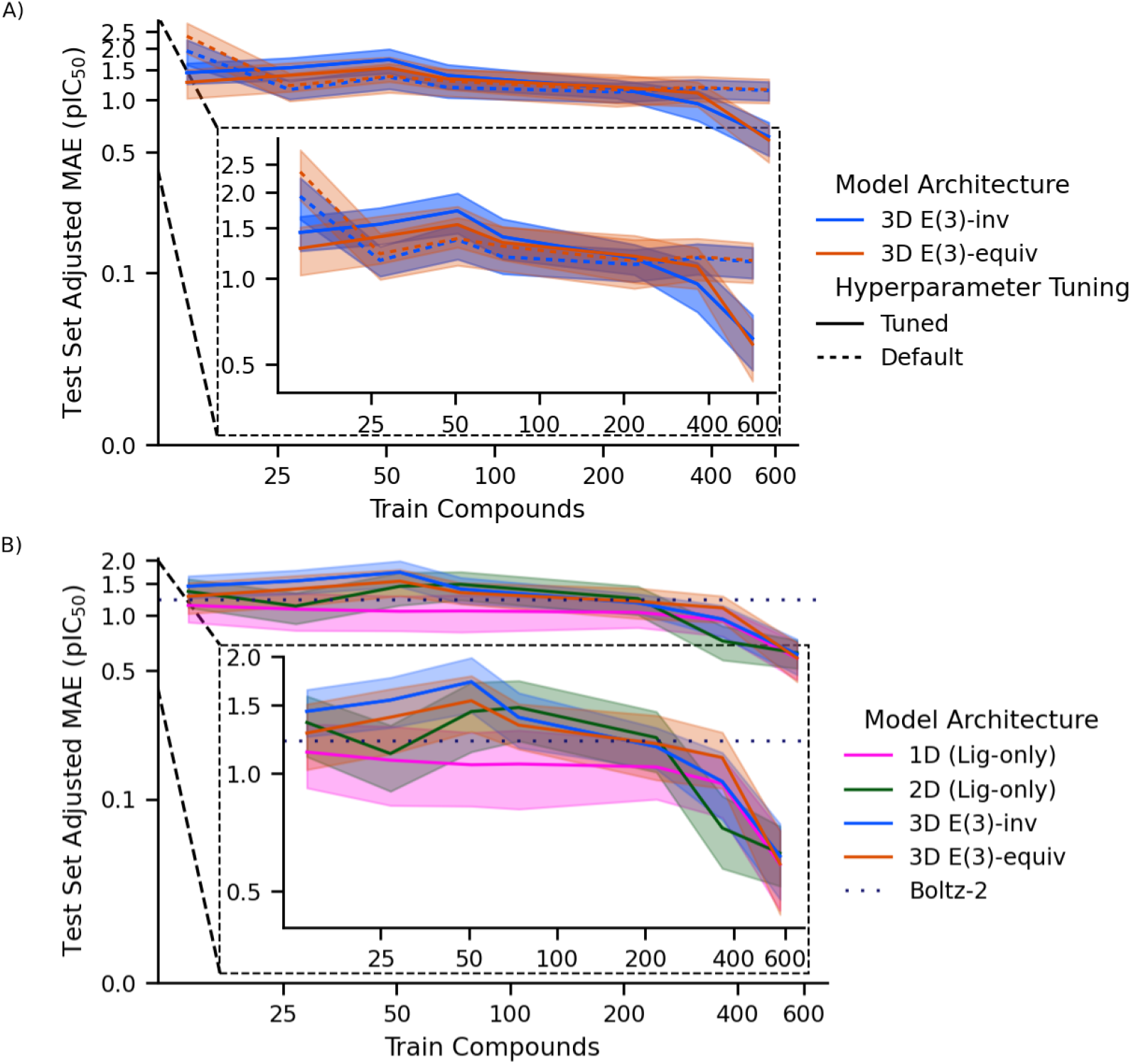
Hyperparameter tuning of structure-based models using PDBBind improves performance on the COVID Moonshot dataset over baseline model construction, enabling performance on par with standard ligand-only models. A. Test set adjusted MAE of our structure-based models after optimizing construction and training. Improvement of tuned models (solid line) is especially visible in the high-data regime, where the default-constructed models (dashed line) do not improve with additional training data. B. Both tuned 3D models (blue and orange) perform comparatively to ligand-only models (green and pink), with indistinguishable performance at the upper end of the available data regime. While all models perform similarly to the Boltz-2 affinity baseline (dotted line) in the low-data regime, once trained on the full available data all four target-specific models outperform Boltz-2.

As before, we are interested in both model performance and model data efficiency, so we evaluate models trained on 1%, 3%, 5%, 10%, 30%, 50%, and 80% of the total available data. To mimic the time-dependent nature of data acquisition in a real-world pipeline, we perform this splitting temporally, with the train set being taken starting from the oldest data, the validation set taken from the following 10% of data, and the most recently generated 10% of the data held as a fixed test set across all train set sizes. Due to the semiquantitative nature of this data, we assess models on their adjusted mean absolute error. This adjustment accounts for predictions that are "semiquantitatively correct", meaning that if an experimental measurement is outside the assay dynamic range, the prediction incurs a loss of 0 if it is outside the assay range in the same manner, i.e. the experimental label and prediction are either both below the assay range of detection or both above the range of detection. The adjusted absolute error for each prediction is calculated as:

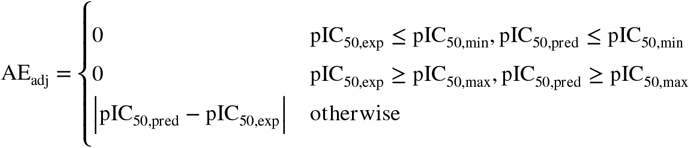

where pIC_50,min_ and pIC_50,max_ represent the minimum and maximum pIC_50_ values detectable by the assay.

As expected, the tuned models for both architectures improve over the default versions (Fig 4A). Although these models perform similarly across most of the range of evaluated dataset sizes, the improvement is especially prominent in the high-data range, where the tuned models are able to improve performance with additional training data while the default-constructed models remain constant in their performance. These results hint at the benefits of our tuning and prediction decomposition approach, showing that these improved models are able to learn more efficiently.

### Hyperparameter-tuned 3D models perform on par with ligand-only models

Having tuned the hyperparameters of our 3D models and verified that their performance has improved as a result, we next compare the performance of these tuned models to 1D and 2D model architecture baselines (Fig 4B). In this experiment, we use the COVID Moonshot dataset prepared as previously described for the 3D models. We construct the inputs for the 1D model using the standard `rdkit` fingerprint, and the 2D model inputs are constructed using `dgl` [80, 88]. Importantly, the compounds used in all splits across models are identical, in order to obtain the most unbiased assessment. As before, we assess model performance across all subsets of the full dataset in order to evaluate overall model performance as well as the degree to which these models’ performance improves with additional data. We also include the performance of the 1D and 2D models in this comparison, as well as the performance of the Boltz-2 affinity predictions, as the baselines against which we compare the performance of the hyperparameter-tuned 3D models (Fig 4B).

All four evaluated models show similar data efficiency trends, with relatively constant model performance across the low-data regime (<200 compounds) (Fig 4B). Beyond the 200 training compounds threshold, all models exhibit significant improvement in performance with additional data. Additionally, all models reach similar final performance, achieving a test set adjusted MAE of ∼0.5 pIC_50_ units. As the performances of all four models are still improving with additional training data at the maximum available amount of data, we hypothesize that the size limitations of the available dataset may be confounding these results to some degree. Future work with larger dataset sizes, such as those currently being generated by the ASAP Discovery Consortium and OpenBind, may help confirm or deny this hypothesis. It is also important to note that, while the 3D models have access to more information in their inputs than the 2D models, the ligand-only models are especially well-suited to this task of training and evaluation on a single target-specific dataset. We find that the additional complexity of the 3D models cancels out any benefit from the extra information, resulting in similar performance to the ligand-only models on this task.

In comparison to the Boltz-2 affinity predictions, while all four models perform similarly to this external baseline in the low-data regime, when trained on sufficient data our evaluated models all outperform the Boltz-2 baseline (Fig 4B). Importantly, the full COVID Moonshot dataset was deposited to the PDB prior to the Boltz-2 training cutoff, meaning that all data in this dataset, including the test set, was present in the Boltz-2 training data [5]. The relative high performance of our evaluated models supports the utility of these less complex, target-specific models, as compared to larger foundation models.

### Pretraining and generalizability tasks highlight efficiency of 3D models

Although the 3D models and 2D baselines performed similarly in our target-specific training and evaluation task, one of the key advantages of structure-based models is their inherent generalizability [43, 89]. We evaluate the ability of these models to generalize to unfamiliar targets on two axes. First, we assess the degree to which pretraining on the PDBBind dataset accelerates fine-tuning on the COVID Moonshot dataset(Fig 5A). We next turn to the task of inference, using a dataset that is fully orthogonal to both the PDBBind and the COVID Moonshot datasets to assess the ability of these models to generalize to previously unseen targets (Fig 5B).

**Figure 5.**
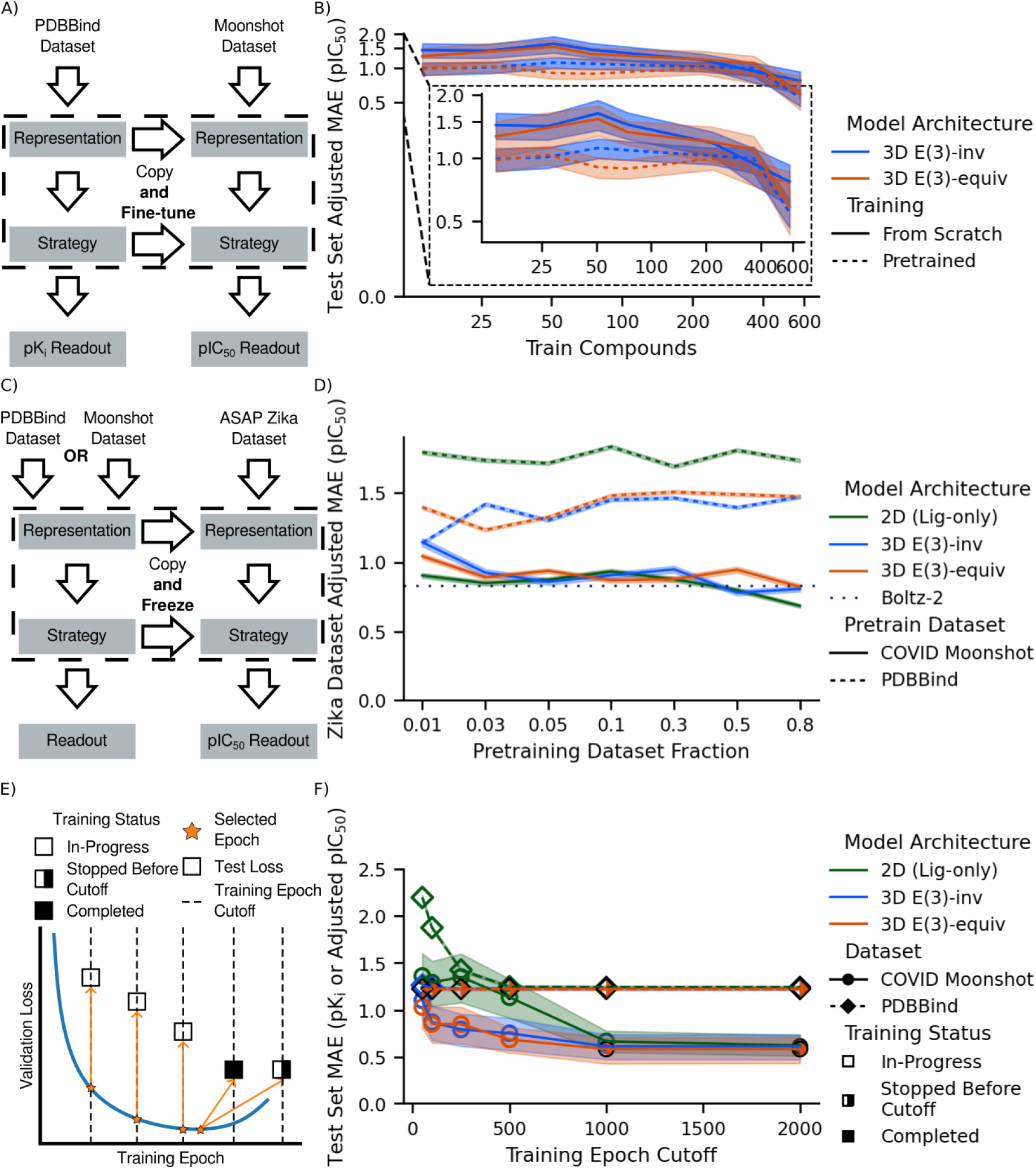
Structure-based models show promise in generalizability, especially in the contexts of pretraining and training efficiency. A. Schematic workflow of model fine-tuning by first training on the PDBBind dataset, using a pK_i_ *Readout*, then copying the learned weights into a new model with a pIC_50_ *Readout*, which is then trained on the COVID Moonshot dataset. B. Performance of structure-based models with and without pretraining (dashed and solid lines, respectively) across a range of train set sizes (x-axis) for both invariant and equivariant models (blue and orange, respectively) C. Schematic workflow of model inference by first training on either the PDBBind dataset or the COVID Moonshot dataset, then using learned model weights to predict on the ASAP Zika dataset. D. Performance of 2D (green) and 3D models (blue and orange) on the inference task across a range of train set sizes, when trained on either the PDBBind (dashed line) or the COVID Moonshot dataset (solid line). E. Workflow to assess training efficiency by limiting the number of training epochs and comparing model performance on the held out test set. Open markers indicate that the minimum validation loss reached up to this training limit is not the lowest validation loss reached across the full pretraining run, half-filled markers indicate that model training was cut off by the wall time limit before this epoch limit, and filled markers indicate that training continued past this epoch cutoff but that validation loss did not further decrease. F. Model performance across different epoch limits of training. This difference is especially marked in the COVID Moonshot dataset (circles, solid lines), where the 3D models (blue and orange) both show steep improvement from 50 epochs of training to 100 epochs, while the 2D model (green) only shows gradual improvement until 1,000 epochs of training. In the high epoch limit of training, there is no difference in model performance between the ligand-only models and structure-based models.

#### Pretraining on the PDBBind dataset accelerates model fine-tuning in the low data regime

To evaluate the ability of these models to generalize from a general binding affinity dataset, we first train the models on the previously described leak-proof splitting of the PDBBind, then further fine-tune the model weights using the COVID Moonshot dataset (Fig 5A). As previously discussed, one of the key features of `mtenn` is its modularity, and the associated ease of pretraining a model on one type of data (K_i_/K_d_ values in this case) and fine-tuning on a different type of data (IC_50_ values in this case). For this pretraining process, we initially train models on the maximal PDBBind dataset split used previously (80% of the total available training set). After training for 24 hours, up to 2,000 epochs, we select the model weights from the epoch with the lowest validation loss to use as the starting weights for training on the COVID Moonshot dataset.

For both the E(3)-invariant and E(3)-equivariant models, we see that pretraining has a positive effect on model performance (Fig 5A), implying that these models are able to generalize knowledge of protein-ligand binding affinities. However, pretraining does not seem to affect the data efficiency of these models, with both models still displaying the same training dynamics with respect to the size of the training set. Although the pretrained models perform better on smaller amounts of data than the trained-from-scratch models, their error similarly does not decrease with more training data up to the critical point ∼200 compounds. This once again hints at the limitations of our dataset [5], as both models are still improving with more data and may show different behavior in higher data regimes.

#### Model inference on an unfamiliar target highlights the generalizability of 3D models

To assess the degree to which the similarity of training and evaluation datasets might be biasing our results (Fig S5), we also turn to new targets to asses model performance and generalizability. Specifically, we turn to the semi-public Zika NS2B-NS3 dataset generated by the AI-driven Structure-enabled Antiviral Platform (ASAP) Discovery Consortium [5, 7, 42]. Similarly to the COVID Moonshot dataset, this Zika dataset contains semiquantitatively labeled binding affinity data, generated for multiple ligands against a single target [7, 21]. Here, we pretrain models as previously described, using either the PDBBind dataset or the COVID Moonshot dataset (Fig 5C). In contrast to the previous fine-tuning analysis, we now freeze all model weights and only perform inference on the Zika dataset compounds. This type of analysis simulates early-stage drug discovery, where there is little to no labeled data, and the goal is to select compounds for initial development. We also vary the size of the pretraining dataset to evaluate the dependence of model generalizability on pretraining dataset size.

The first comparison we investigate is between models within a given pretraining dataset (Fig 5D). For models pretrained on the COVID Moonshot dataset, there is little difference in performance between models across all pretrain dataset sizes. These results hint that we are once again reaching the limits of what we can assess with the dataset, highlighting the difficulty in working with these minimal target-specific datasets. Despite these dataset limits, our models pretrained on the COVID Moonshot dataset all perform similarly to the Boltz-2 baseline, once again highlighting the strength of these target-specific models, even when the inference target differs from the pretraining target.

For models pretrained on the PDBBind dataset, there is a clear gap in model performance on this inference task across all pretrain dataset sizes. This generalization gap implies that the structure-based models are better at generalizing from this target-nonspecific dataset to a target-specific dataset.

Of note in these results is the difference in data label composition between the PDBBind pretraining dataset and the Zika inference dataset: although the Zika dataset is almost 50% comprised of semiquantitatively labeled data points, the PDBBind dataset does not contain any semiquantitative labels. This is a true out-of-distribution generalization, and supports the hypothesis that these 3D models are able to learn information that allows them to predict whether a protein-ligand complex will bind. It becomes clear that these semiquantitative data points present a large source of this generalization gap and are a significant difficulty for all three models, as the generalization gap shrinks and performance for all models improves greatly when the MAE is computed only on in-range data points (Fig S6).

Comparing model performance between pretraining datasets paints a clear pictures of the utility of pretraining on target-specific datasets. For all three models, performance on this inference task is significantly improved by pretraining on the COVID Moonshot dataset as compared to pretraining on the PDB-Bind dataset (Fig 5D). This difference is emphasized by the consideration of the number of compounds in each pretraining dataset. Although we plot model performance against the fraction of the dataset used in pretraining for the sake of comparison, as previously discussed, the PDBBind dataset is orders of magnitude larger than the COVID Moonshot dataset. Despite this difference in absolute size of the pretraining dataset, all three models are able to learn more from the Moonshot dataset. Further, the targets in these two datasets are different proteins from different viral families, implying that a large portion of the benefit from these datasets come from the similarity in data types (i.e. semiquantitively labeled affinity data), as well as simply the fact that these datasets are target-specific.

Another important consideration with these results is the chemical similarity of the molecules in the pretraining dataset set and the molecules in the inference Zika dataset. The distributions of Tanimoto similarities using the rdkit fingerprint show that the chemical space is quite similar between both pretraining datasets and the inference dataset (Fig S5). Pairing the chemical similarity distributions with binding affinity similarities clarifies this relationship further. Although the compounds in the inference dataset are chemically similar to the compounds in both pretraining datasets, the affinity labels for chemically similar molecules differ much more in the PDBBind dataset than in the COVID Moonshot dataset. This difference helps explain the drastic difference in performance for the 2D model between the two pretrain datasets: as the ligand-only model can only learn a mapping between chemical input and affinity output, it expects inputs that are chemically similar to have similar binding affinities. Structure-based models are able to synthesize information about the target in the input, and therefore suffer less from this difference in binding affinity labels for chemically similar compounds. Although complicated by the limits of the available data, the results in this section highlight the potential benefits of using structure-based models for these generalizability tasks.

### Limiting training epochs shows the efficiency of training structure-based models

In previous experiments, all models have been allowed to train for a maximum of 24 hours, up to 2,000 epochs. Although this time limit allows sufficient time for models to train on the COVID Moonshot dataset (Fig S7-S9), due to the size of the dataset, some of the models were unable to finish training on the PDBBind (Fig S10-S12). This training limit is arbitrarily selected, but allows an even comparison of models from a practical standpoint, as wall time is likely to ultimately be the limiting factor in model training, as opposed to the number of training epochs. However, this wall time-based limit inherently puts the 3D models at a disadvantage compared to the ligand-only models, given the extra mathematical complexity. In this analysis, we compare model training efficiency as a function of training epochs rather than as a function of dataset size, as in previous experiments.

To ensure models are compared fairly in terms of training epochs, we limit each model to training up to 50, 100, 250, 500, 1,000, or 2,000 epochs, with the same wall time cutoff of 24 hours (Fig 5E). As before, for each training cutoff, we select the epoch with the lowest validation loss, now restricting the available epochs to those before the cutoff, and compare on the test set loss at the selected epoch. We also annotate the training status at the selected epoch: for training epoch cutoffs where the overall lowest validation loss was not reached, training is classified as "In-Progress"; for training epoch cutoffs where training stopped before the cutoff, due to the wall time limit, training is classified as "Stopped Before Cutoff"; for training epoch cutoffs where the overall lowest validation loss has been reached and training continues at least up to the cutoff, training is classified as "Completed". The distinction between "Stopped Before Cutoff" and "Completed" is important, as the prior label indicates that the model may still be improving in its validation loss, while the latter label indicates that we know that the model does not achieve a lower validation loss before the epoch cutoff. These labels are applied based on a consensus of all model seed replicates: the "In-Progress" label is applied if any of the replicates did not reach the overall minimum validation loss; the "Completed" label is applied if all of the replicates reached the overall minimum validation loss and none of the replicates had their training stopped before the epoch cutoff; the "Stopped Before Cutoff" labels is applied if all of the replicates reached the overall minimum validation loss and any of the replicates had their training stopped before the epoch cutoff.

This analysis highlights the strength of these 3D models’ training efficiency. In both datasets, the structure-based models outperform the ligand-only model at the low-epoch limit of training. In the case of the PDB-Bind dataset, there is a large initial gap in performance that is closed by the 2D model after ∼500 epochs of training. There is a similar trend with the COVID Moonshot dataset, where the structure-based models outperform the ligand-only model at lower epochs of training but all models perform similarly at higher epochs of training. Although this effect occurs at a later epoch in the Moonshot dataset, this is likely due to the smaller dataset causing the 2D model to learn more slowly. These patterns support the hypothesis that the inclusion of physical constraints in the structure-based models improves their training efficiency, allowing them to learn more efficiently than the ligand-only model.

One caveat to note is that both 3D models were cut off in training due to the wall clock time limit after 50-100 epochs of training (denoted by the half-filled markers). Although this limitation does not affect the conclusion that the structure-based models are able to learn in fewer epochs than the baseline ligand-only model, it is possible that the 3D models’ performance will continue to improve with more training epochs. The compute time requirements for training the structure-based models on the full PDBBind dataset prevent us from being able to fully train these models to 2,000 epochs, however an extended analysis with a smaller subset of the PDBBind (Fig S13) shows that the performance of the 3D models does not improve with more training, providing further evidence that these models quickly reach their maximal performance. While this analysis may not be practically useful at present, the results show that structure-based models learn more efficiently than ligand-only models, highlighting the importance of their physical biases. As hardware improves and model architectures are further optimized, the gap in wall time efficiency may also shrink, leading to real, practical benefits for structure-based models.

## Conclusion

In this work, we characterize structure-based models for protein-ligand binding affinity, assessing performance improvements from including physics-based priors in model construction. To assist in these analyses and facilitate fine-tuning during model construction, we present `mtenn`, a Python package that formalizes our decomposition of the affinity prediction task into modular components to encode substructures of the protein and to combine these learned embeddings. Using this decomposition, we evaluate different levels of priors at both the structure embedding (*Representation*) and embedding combination (*Strategy*) steps. In the structure embedding stage, we compare E(3)-invariant and E(3)-equivariant models to measure the improvement enabled by using these physical constraints. In the embedding combination stage, we compare a *Strategy* based on the definition of the binding affinity with one that instead allows a neural network to learn how different substructures of the protein-ligand complex interact.

We first perform hyperparameter tuning using a leak-proof splitting of the PDBBind [81], and show that this tuning on a general binding affinity dataset improves model performance on the target-specific drug discovery dataset of the COVID Moonshot dataset [54]. With proper tuning, structure-based models perform on par with ligand-only models. We find that while inclusion of physics-based priors at the stage of combining learned representations has a positive effect on model performance, there is no benefit to using the theoretically more expressive E(3)-equivariant model as compared to the E(3)-invariant model. Although these results are promising, there are limitations in working with the COVID Moonshot dataset. This dataset is one of, if not the largest publicly available target-specific binding affinity dataset. Despite the impressive resource of the Moonshot dataset, we remain limited by the lack of data relative to datasets like the PDB-Bind, highlighting the need to continue investing time and money in generating these public datasets [5].

In addition to their performance on the plain binding affinity prediction, we also assess the models’ ability to generalize to unfamiliar data. We find that all models are able to generalize similarly well when pretrained on the PDBBind dataset and subsequently fine-tuned on the COVID Moonshot dataset, although as previously mentioned, there is noise due to the limitations of the Moonshot dataset. Turning to a pure out-of-distribution inference task, structure-based models are better able to generalize from the PDBBind dataset to the Zika dataset, although there is no difference in performance between all models when pretrained on the Moonshot dataset. Finally, when considering efficiency in terms of training epochs, we show that structure-based models are more efficient than the ligand-only model in training on both the PDBBind dataset and the COVID Moonshot dataset.

Throughout this paper, we repeatedly encounter limitations in our analysis due to available data sources. Despite the success of the COVID Moonshot project, and the continued efforts of the ASAP Discovery Consortium, the availability of public drug discovery campaign data remains low in comparison to the amount of publicly available binding affinity data. Forward-thinking drug discovery companies are already working to alleviate the bottlenecks of limited data regimes by turning to federated learning, using companies like Apheris to allow them to benefit from other companies’ proprietary data without needing to disclose their own data [90].

While this model works well in the private sphere, public depositions of drug discovery campaign data remain uncommon. Recent dataset and model releases from the OpenBind consortium emphasize the benefits of additional datasets in new contexts for the field [29, 91, 92]. Furthermore, the deployment of blind challenges enables us to generate continuous rigorous benchmarks to assess state-of-the-art improvement [21]. We believe that the resulting inequities in model performance across sectors are, at least in part, due to the access disparities we note above. Our work above outlines methods to improve model performance using publicly available datasets, but increasing accessible data is another sure method of model improvement that will open new avenues of model development.

Looking to the future, this work demonstrates the utility of structure-based machine learning models for binding affinity prediction in the context of drug discovery; the benefits are most visible during the earlier stages of a design campaign where training data is limited and generalizability is of utmost importance. We expect the bottleneck in model performance to shift towards the amount of training a model is able to accomplish. As hardware improves and model architectures become more advanced and efficient, and structure-based models require less wall time per epoch, the efficiency of these physics-based architectures will continue to improve performance.

## Detailed Methods

### The Leak-Proof PDBBind Dataset

The structures used for the PDBBind dataset were generated from the v2020 version of the dataset, [81] downloaded from the PDBBind website. To combine the protein and ligand structures that are stored in the PDBBind, we used the `drugforge` package, combining the protein structures stored in the PDB files with the ligand structures stored in the MOL2 files [42]. We used the MOL2 files rather than the SDF files for this data generation as we found inconsistencies in bond order annotation in the SDF files. The train, validation, and test splits were subsequently generated using the `new_split` column in the `dataset/LP_PDBBind.csv` file downloaded from the leak-proof PDBBind GitHub [https://github.com/THGLab/LP-PDBBind/tree/master] on September 19, 2024 [93].

### The COVID Moonshot Dataset

The ligand data for the COVID Moonshot Dataset was downloaded from GitHub (https://github.com/foldingathome/covid-moonshot), and filtered to only include achiral and enantiopure molecules using drugforge-data [54].

The protein structures used to generated the COVID Moonshot dataset were downloaded from the Diamond Light Source app Fragalysis [94] on October 15, 2024, using their API. The structures were then prepped using the Spruce Modeling Toolkit from OpenEye, accessed via the `drugforge-modeling` package. Docked structures were then generated with OpenEye POSIT docking, accessed via `drugforge-docking`, using the prepped crystal structures and ligand SMILES strings as input [1]. To determine to which reference crystal structure each ligand was docked, a maximum common substructure (MCS) search was performed for each query ligand, and the crystal structure whose ligand shared an MCS with the query ligand was selected as the reference. Within the docking protocol, we ran an OMEGA dense search for each ligand, returning a maximum of 50 poses [87]. For the single-pose analysis in the paper, the docking pose with the top POSIT score was selected as the complex structure for that ligand. For the multi-pose analysis, all returned ligand poses were filtered down to poses that were unique up to 2Å. This filtering was performed by iterating through the list of poses, sorted by decreasing POSIT score, and keeping all poses that had > 2Å RMSD to all previously selected poses.

### Subsampling of the Leak-Proof PDBBind

For the data efficiency experiments using the PDBBind dataset, we randomly subsampled the leak-proof splits using fixed random seeds. For the training and validation sets, we randomly selected 1%, 3%, 5%, 10%, 30%, 50%, and 80% of the PDB IDs in each split given by the leak-proof splitting, using fixed random seeds of 1 and 2. For all experiments using the PDBBind, we used the full test set as given by the leak-proof splitting, to ensure appropriate comparisons across experiments.

### Subsampling of the COVID Moonshot Dataset

For the data efficiency experiments using the COVID Moonshot dataset, we subsampled the dataset temporally, using 1%, 3%, 5%, 10%, 30%, 50%, and 80% of the total available compounds in the training set. Because this subsampling was done deterministically, there is only one replicate for each training set size. To compute each split, we add compounds by ideation date, ensuring that all compounds from the same date are in the same set. As with the PDBBind subsampling, the test set is fixed to ensure comparisons are equivalent. In this case, the test set is fixed as the most recent 10% of compounds. The training set is calculated by rolling the cutoff date across all sorted ideation dates in the dataset and adding all compounds from that date to the train set, until the appropriate training set size is reached. The validation set is calculated in the same way, starting with the first date in the dataset after the end of the training set. The size of the validation set is also fixed at 10% of compounds for all training set sizes.

### Confidence Interval Calculation

For all experiments, confidence intervals were calculated using the `scipy.stats.bootstrap` function with the appropriate function. We bootstrapped all metrics with 9,999 resamples, using the `basic` method with a `two-sided` alternative. The code for performing this bootstrapping is available in `drugforge.ml.analysis.calc_stats`.

While rigorous statistical analyses are typically carried out using cross-validation (CV) [85], standard CV approaches are not well suited to time-labeled data. Methods that are suited to temporal data employ a sliding window approach, where the initial fold contains data with the earliest time labels, and each subsequent fold contains data from the previous fold as well as additional future data [95]. This sliding window approach is identical to the subsampling approach we employ in our data efficiency experiments, making these two approaches mutually exclusive. Due to the difficulty of performing proper cross validation with temporally split data, we compare experimental conditions based on the overlap of these confidence intervals.

### Hyperparameter Tuning on the PDBBind

The protein and ligand structures were downloaded from PDBBind, and the complex structures were generated by combining the protein and ligand structures using the `drugforge` software package. In order to reduce the computational time needed for each of the experiments in this section, we use 50% of the available training and validation sets to perform our hyperparameter tuning experiments.

Rather than testing every possible combination of hyperparameters (Table S1), we limit the total number of training runs required by tuning sequentially, selecting the best option for each model/*Strategy* pair at each step and fixing it for subsequent tuning experiments (Fig 3A). At each tuning step, we run experiments on the E(3)-invariant and E(3)-equivariant models, and the Delta and Concat *Strategy*, and for each experiment, we run all models on both subsamplings of the 50% dataset, and with two fixed random seeds for generating initial model weights. At this step of hyperparameter tuning, we vary 5 different aspects of model construction and training.

We first test three different loss functions to tune how the training process accounts for outliers in model predictions: an L1 (mean absolute error) loss, which is more robust to outliers but overly penalizes models when the error is small; an L2 (mean squared error) loss, which slows training when the error is small, but greatly increases the loss for larger errors; and a Smooth L1 loss, which combines these two by imposing a squared loss term for small errors and a linear loss term for larger errors (Fig S1). Our analysis shows minimal differences between the tested loss functions, so we select the Smooth L1 loss as the loss function to be used in subsequent training runs for all model/*Strategy* pairs, in order to balance the strengths of the L1 and L2 loss functions.

We next tune the minibatch size and learning rate used in training. These hyperparameters are intrinsically linked, as the amount of data seen per model step informs how large of a step should be taken, so we perform a grid search over the selected space for these values. We range the learning rate from 10^−5^ to 10^−3^ and the minibatch size from 1 sample per optimizer step to the entire train set. The full list of values, as well as the selected values for each model/*Strategy* pair are shown in Table S1 and Fig S2.

We next evaluate the models’ reliance on memorizing the order of the training set by randomizing the order in which the data is shown to the model each epoch (Fig S3). We find no significant difference in model performance between shuffling and not shuffling the train set. We opt to randomize the data order for each epoch in all experiments going forward to improve model robustness.

We also tune the models’ memorization of atomic coordinates by applying random jittering to all atom positions (Fig S4). For each atom in each structure in the train set, a random value is drawn from a normal distribution with a mean of 0 and varying standard deviation for each coordinate. This random noise is added to the atoms’ position vectors, and is drawn independently for each epoch. We compare 5 different values for the standard deviation of the noise distribution (0.01Å, 0.05Å, 0.1Å, 0.5Å, 1Å) as well as passing the original fixed structures (no jittering). We also evaluate noise distributions with standard deviations of 10Å and 100Å amounts as a control for this experiment, as we expect this degree of positional distortion to seriously impact the model’s ability to learn. As expected, a noise distribution with a standard deviation of 100Å significantly worsens model performance. The 10Å distribution also has a significant negative effect on model performance for both models with the Delta *Strategy*, but a negligible effect on both models with the Concat *Strategy*, suggesting that the Concat *Strategy* may be more robust to variations in the input structures. The effect sizes for the less extreme noise distributions are minimal, but based on the results we select a standard deviation of 0.1Å for E(3)-invariant/Delta, 1Å for E(3)-invariant/Concat, 0.5Å for E(3)-equivariant/Delta, and 0.5Å for E(3)-equivariant/Concat.

## Data and Code Availability

The `drugforge` toolkit code is open-source and available on GitHub [https://github.com/choderalab/drugforge] under a permissive MIT License. The `mtenn` framework code is similarly available on GitHub [https://github.com/choderalab/mtenn] under a permissive MIT License. Zika virus inhibitors from the ASAP Discovery Consortium are available through the consortium portal [https://asapdiscovery.org]. A fully interactive discovery pipeline that points to all experimental outputs of ASAP Discovery is located at the Consortium’s web site [7]. The complete scripts to training models, running pipelines, analyzing data, and generating figures for this paper are available on GitHub [https://github.com/choderalab/physical-biases-paper]. Our parsed results from the Moonshot and ASAP datasets, alongside complete data input and output files needed to regenerate this work are available on Zenodo [https://doi.org/10.5281/zenodo.22050744].

## Acknowledgments

We are grateful to the ASAP Discovery Consortium [http://asapdiscovery.org] and its numerous talented scientists for their contributions to data generation, scientific motivation, and many thoughtful discussions in the formulation of this work. We thank Jenke Scheen and Ed Griffen for helpful discussions during the preparation of this manuscript. We also thank Iván Pulido, Chris Iacovella, and Mike Henry for help with the `drugforge` package. This work used resources from the High-Performance Computing Group at Memorial Sloan Kettering Cancer Center. The authors are grateful to the MSKCC DigITs and HPC team, especially Jamie Cheong, Lohit Valleru, and Monica Chakradeo for their assistance with high-performance computing resources.

## Disclaimer

The content is solely the responsibility of the authors and does not necessarily represent the official views of the National Institutes of Health.

## Author Contributions

Conceptualization: BK, JDC; Methodology: BK, SS; Software: BK, AMP, HMO; Formal analysis: BK, SS; Investigation: BK, SS ; Writing–Original Draft: BK, SS; Writing–review & editing: BK, SS, HMO; Funding Acquisition: JDC; Resources: JDC; Supervision: SS, JDC.

