## Supporting Information and Figures for "Physical priors improve performance of structure-based binding affinity models"

### Supplementary Information for: Physical priors improve performance of structure-based binding affinity models

| Experiment | Model | Strategy | Loss Function | Data Order Randomization | Minibatch Size | Learning Rate | Positional Jittering |
|---|---|---|---|---|---|---|---|
| **Loss Function (1)** | E(3)-inv<br>E(3)-equiv | Delta<br>Concat | Squared error | No | 1 | 1e-4 | No |
| **Minibatch / Learning Rate (2)** | E(3)-inv<br>E(3)-equiv | Delta<br>Concat | Choose 1 based on exp 1 | No | 1<br>10<br>50<br>100<br>500<br>1,000<br>5,000<br>-1 | 1e-3<br>5e-4<br>1e-4<br>5e-5<br>1e-5 | No |
| **Data Ordering (3)** | E(3)-inv<br>E(3)-equiv | Delta<br>Concat | Choose 1 based on exp 1 | Yes<br>No | Choose 1 per Model/Strategy comb based on exp 2 | Choose 1 per Model/Strategy comb based on exp 2 | No |
| **Jittering (4)** | E(3)-inv<br>E(3)-equiv | Delta<br>Concat | Choose 1 based on exp 1 | Choose 1 based on exp 3 | Choose 1 per Model/Strategy comb based on exp 2 | Choose 1 per Model/Strategy comb based on exp 2 | 0.01 Å noise<br>0.05 Å noise<br>0.1 Å noise<br>0.05 Å noise<br>1 Å noise<br>10 Å noise<br>100 Å noise |

**Table S1. We tune structure-based model hyperparameters using a sequential grid search.**
In each tuning experiment, we perform all possible combinations of the listed parameters (the Cartesian product of all cells in the row). We limit the total number of tuning experiments needed by performing tuning sequentially, fixing the highest-performing parameters in each experiment for all subsequent experiments.

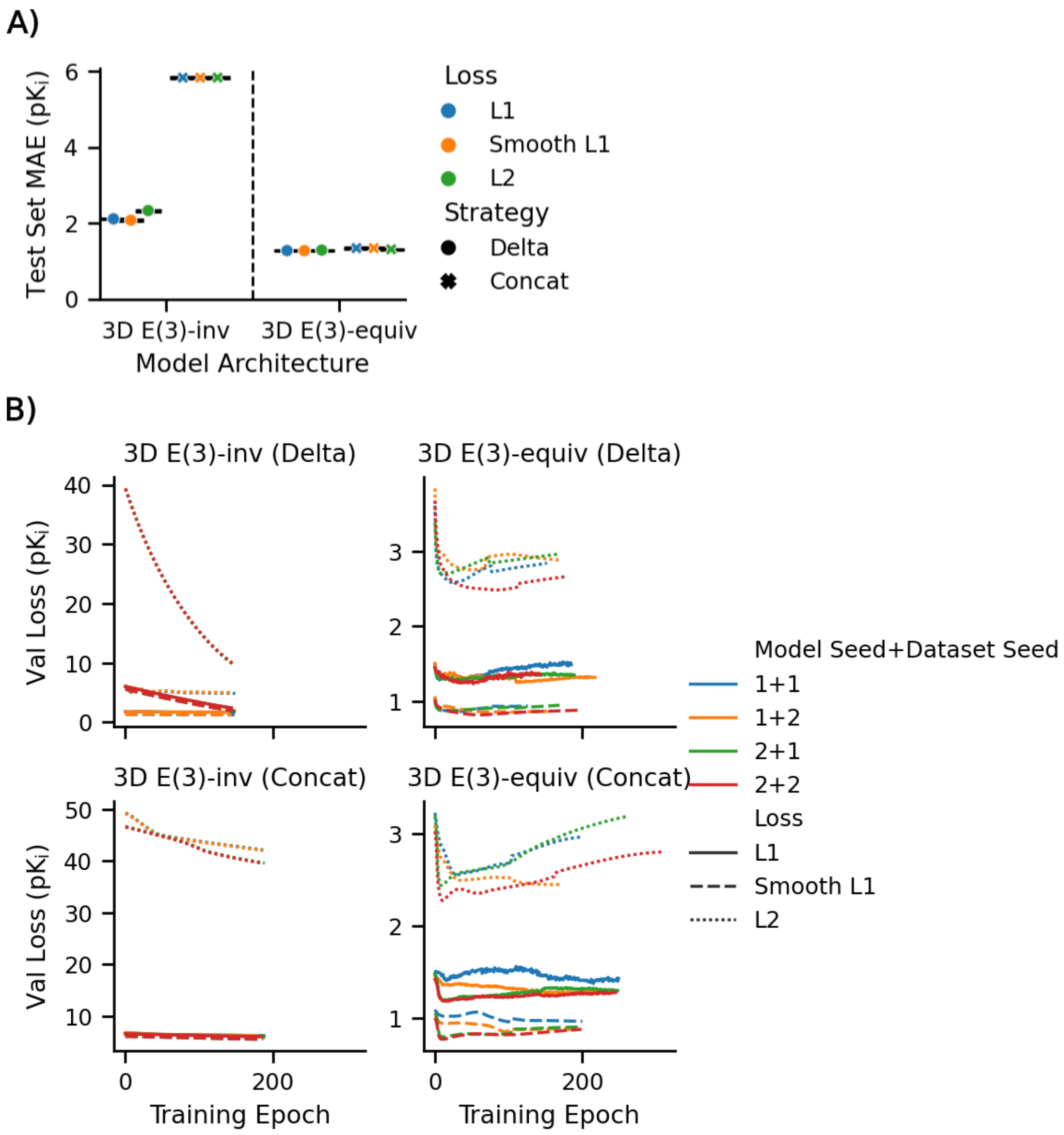


**Figure S1. There are minimal differences in loss function performance.**
A. Test set mean absolute error for all Model/*Strategy* combinations, using each of the three tested loss functions. B. Validation loss curves during training for results in (A).

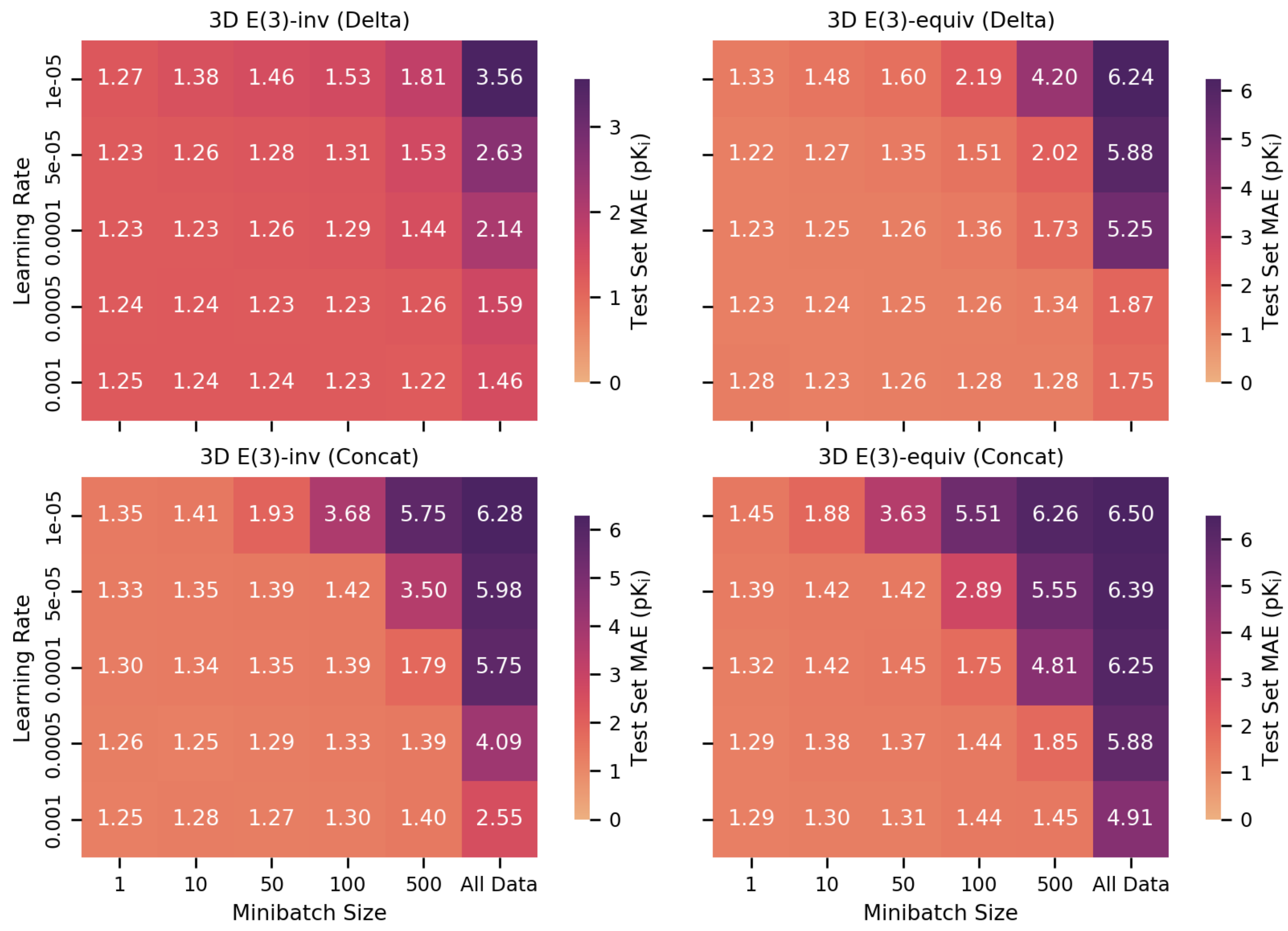


**Figure S2. PDBBind test set mean absolute error for different combinations of learning rate and minibatch size.**

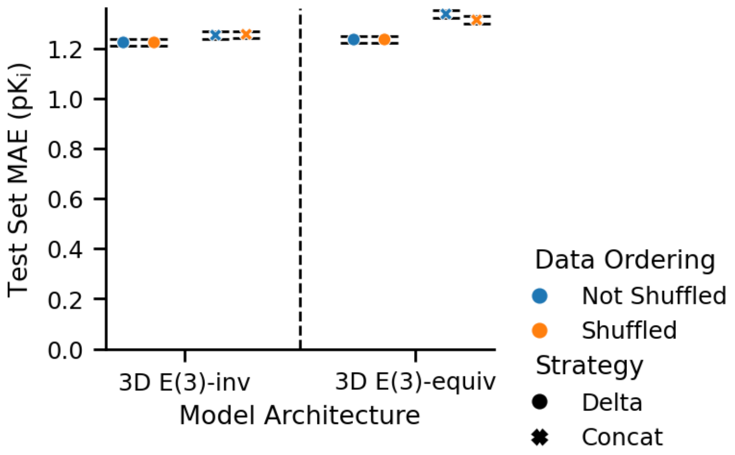


**Figure S3. PDBBind test set mean absolute error when the order of the training set is the same or randomized each training epoch.** Test set MAE is shown for both shuffled and non-shuffled data orderings (orange and blue respectively), and Delta and Concat *Strategy* options (circle and X symbols, respectively)

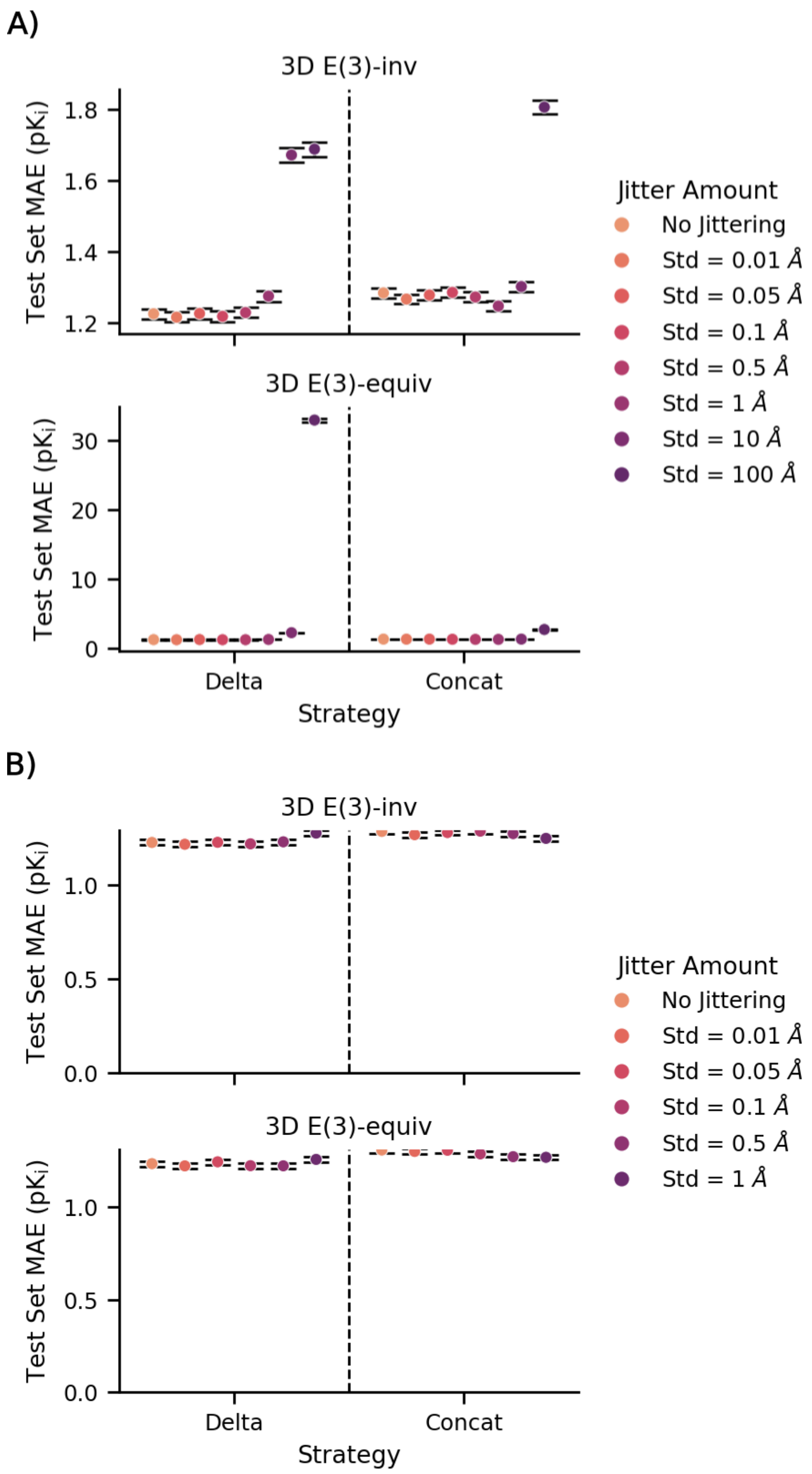


**Figure S4. PDBBind test set mean absolute error with different amounts of positional jittering applied to the train set input coordinates.** Standard deviation of the noise, representing the degree of jittering applied to the dataset, in Å is noted in the legend. Results are presented with (A) and without (B) controls of 10 Å and 100 Å of noise.

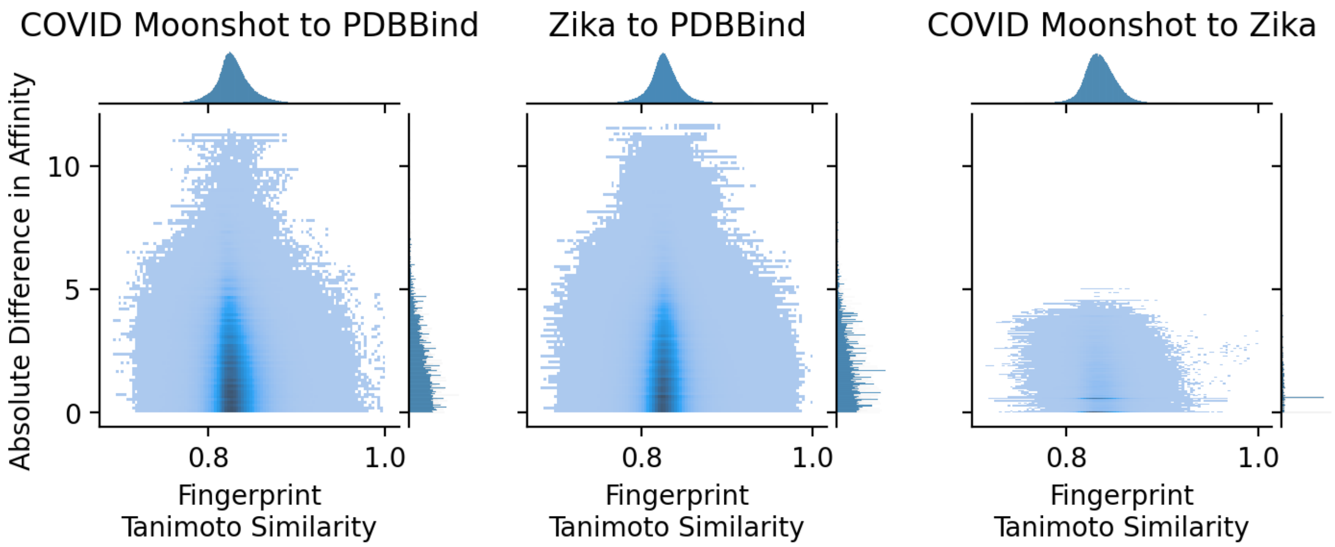


**Figure S5. 2D histogram of Tanimoto similarity using the rdkit fingerprint and binding affinity similarity for all pairwise comparisons between datasets.** All three datasets contain chemically similarly compounds, but both target-specific datasets differ significantly from the PDBBind in terms of affinity similarity. The two target-specific datasets are, however, quite similar to each other in terms of binding affinities.

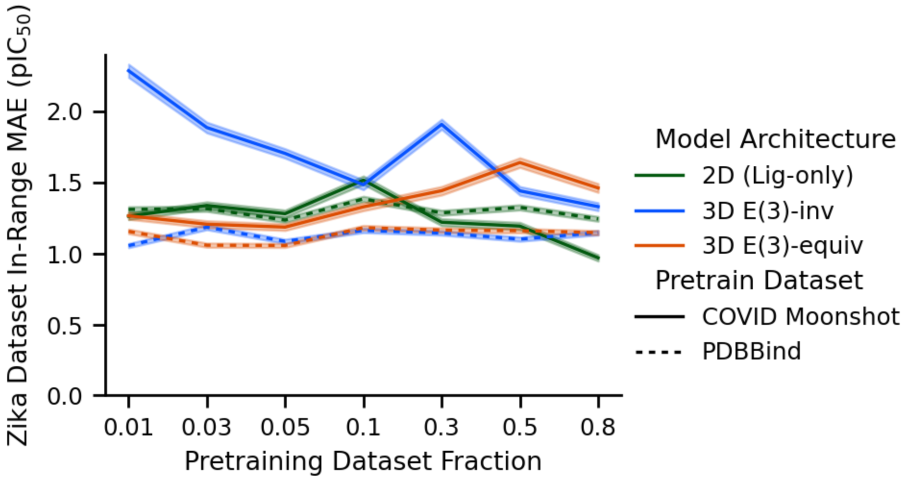


**Figure S6. Zika inference mean absolute error for compounds with in-range affinities only.** In-range MAE for all models, trained using either the COVID Moonshot dataset (solid lines) or PDBBind (dashed lines). Both training datasets are similar, showing the effects of the out-of-range data points on model performance.

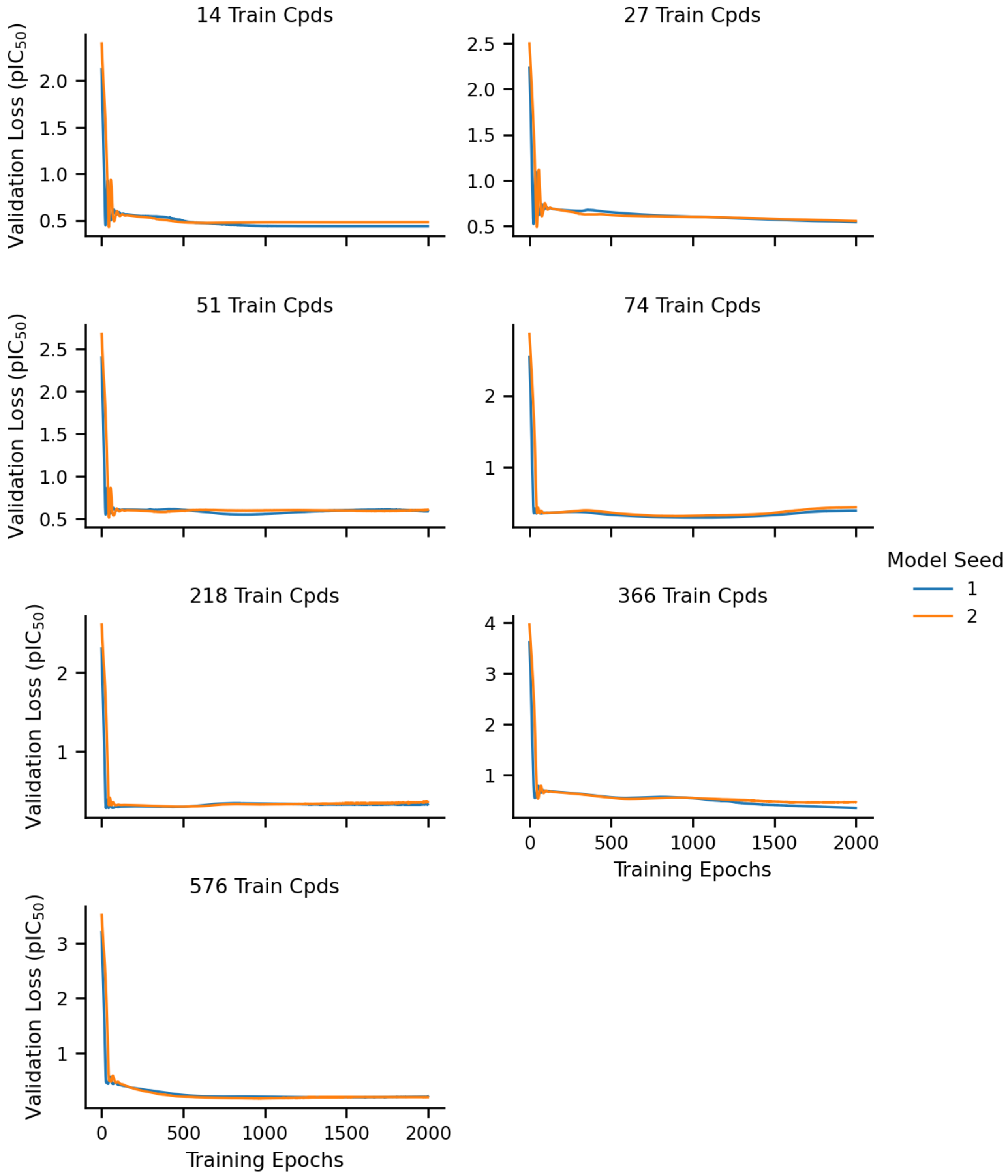


**Figure S7. Training curves for the 2D ligand-only model on the COVID Moonshot dataset.** Validation loss is shown for differing numbers of compounds using two different seeds for generating initial model weights (legend). All replicates appear to have completed training, as validation loss is not decreasing at any dataset size.

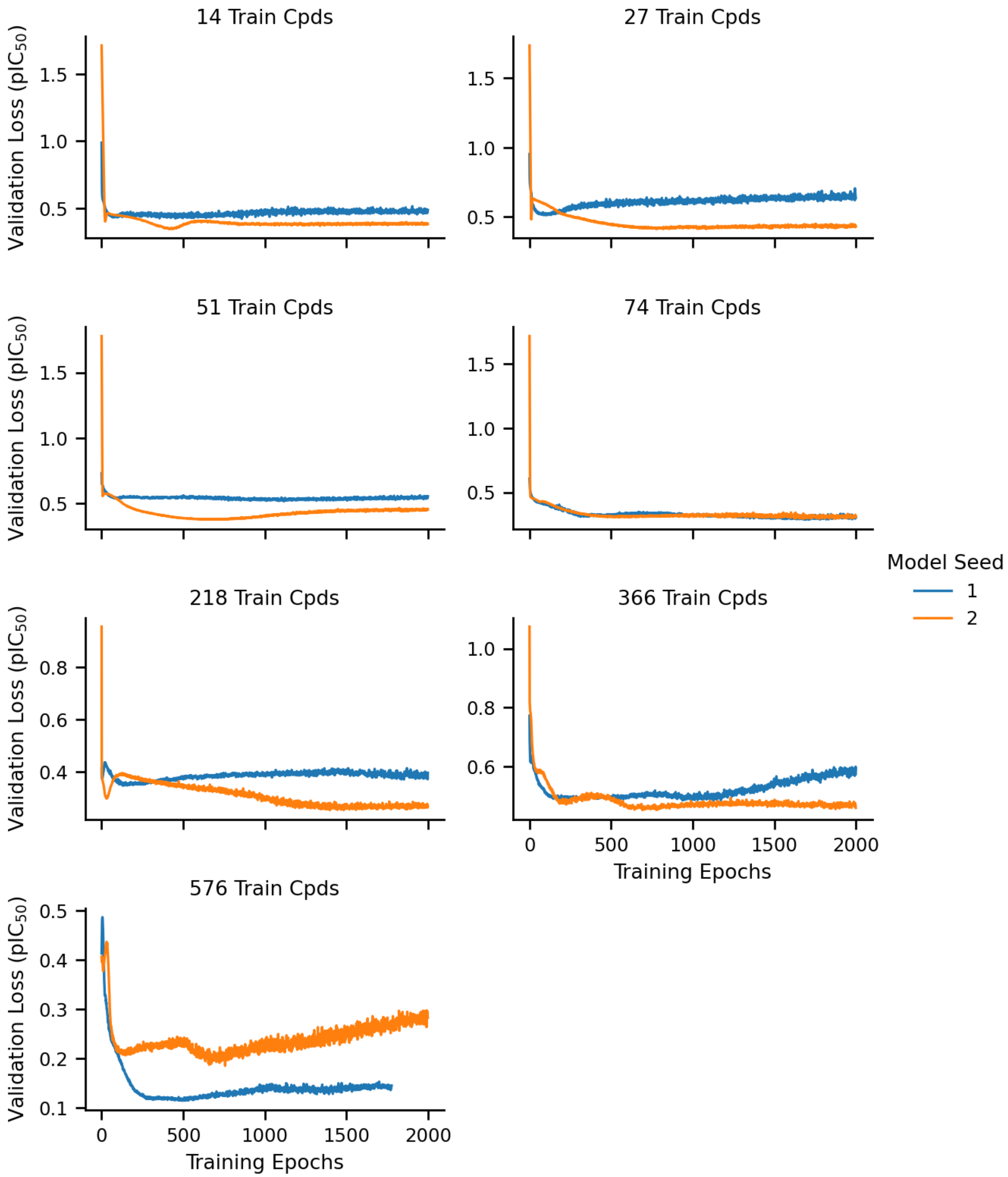


**Figure S8. Training curves for training the 3D E(3)-invariant model on the COVID Moonshot dataset.** All replicates appear to have completed training, as validation loss is not decreasing at any dataset size.

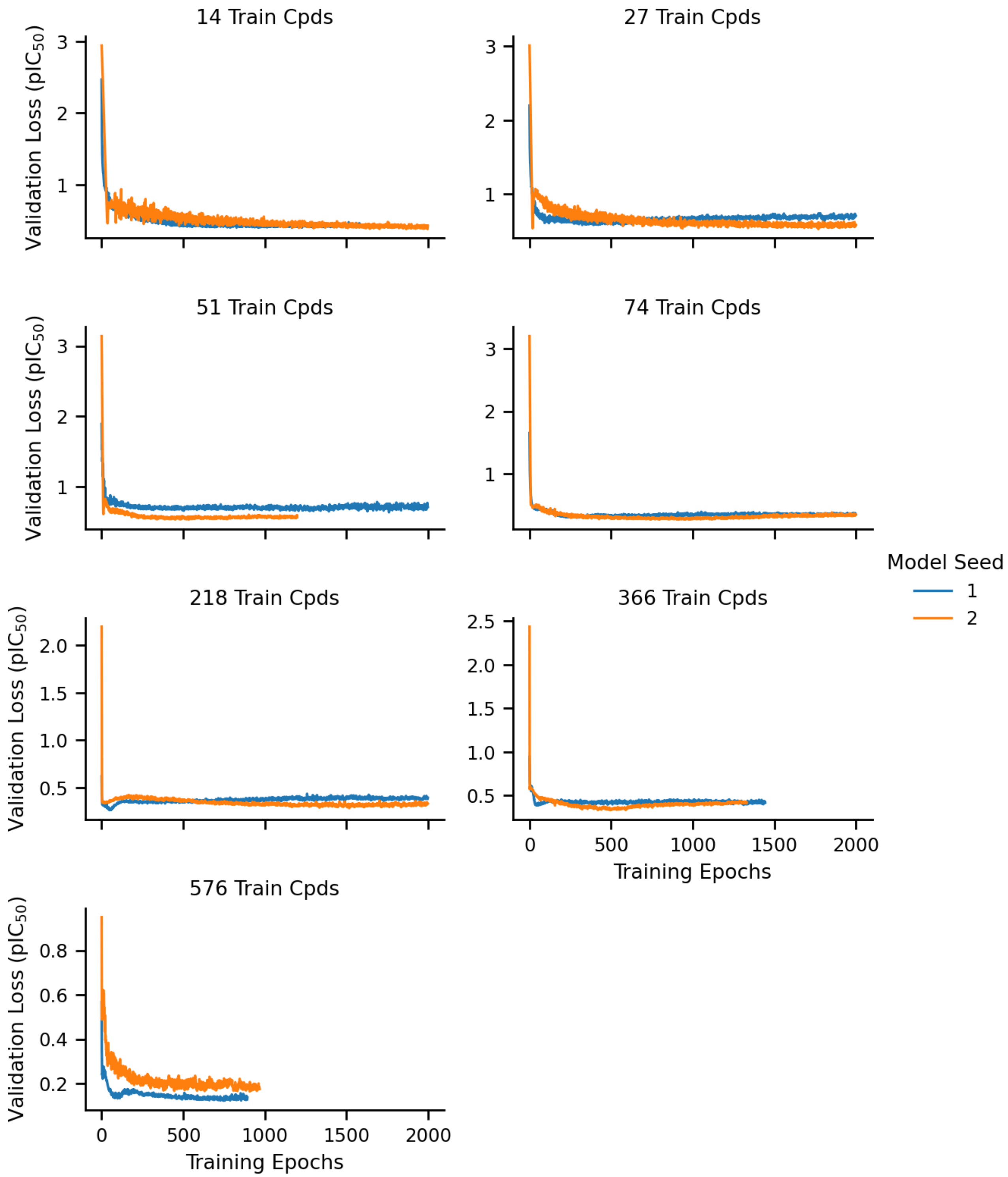


**Figure S9. Training curves for training the 3D E(3)-equivariant model on the COVID Moonshot dataset.** All replicates appear to have completed training, as validation loss is not decreasing at any dataset size.

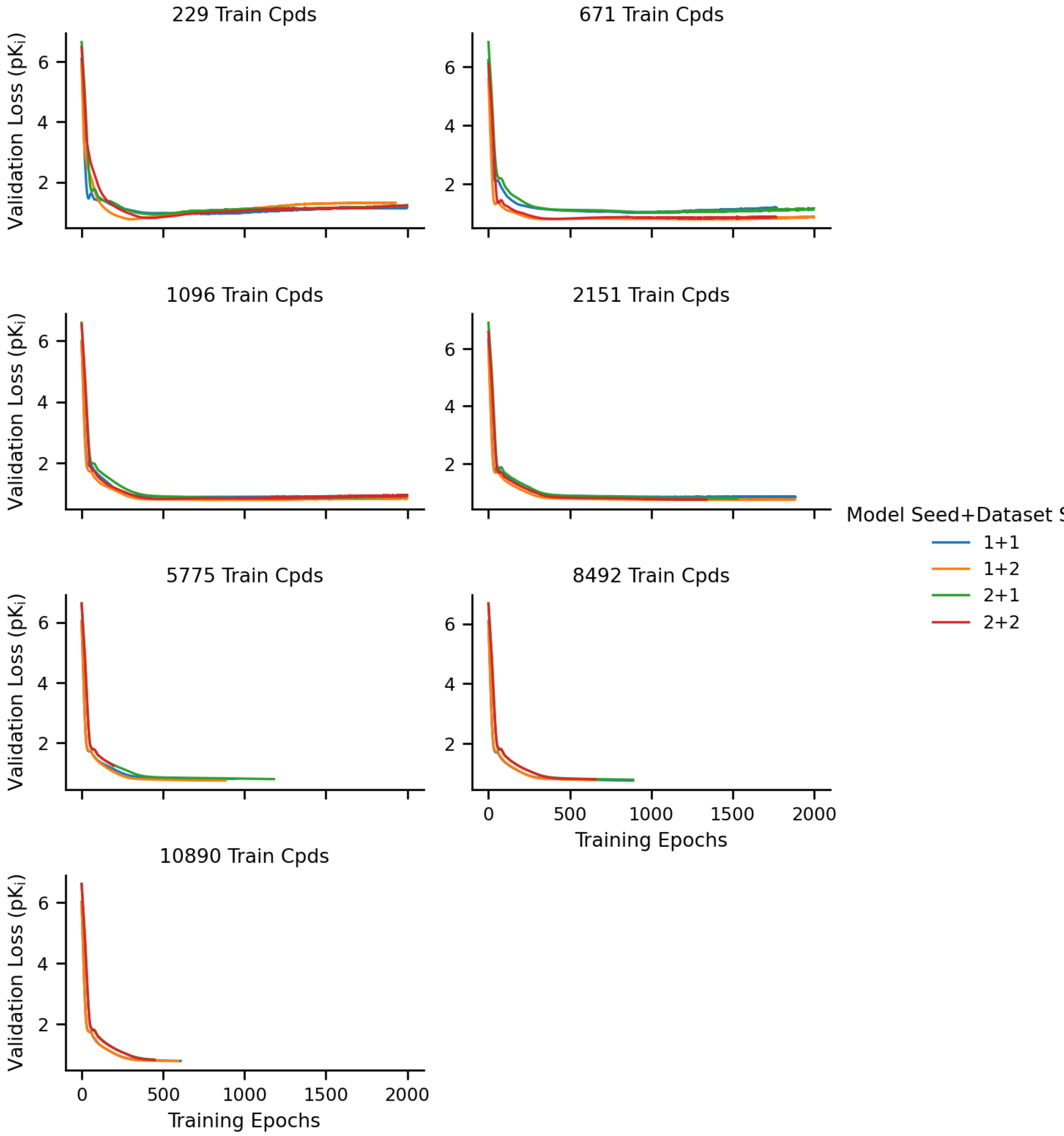


**Figure S10. Training curves for training the 2D ligand-only model on the PDBBind dataset.** Replicates across smaller train set sizes appear to have completed training, as validation loss is not decreasing, while runs with larger train set sizes may still be training.

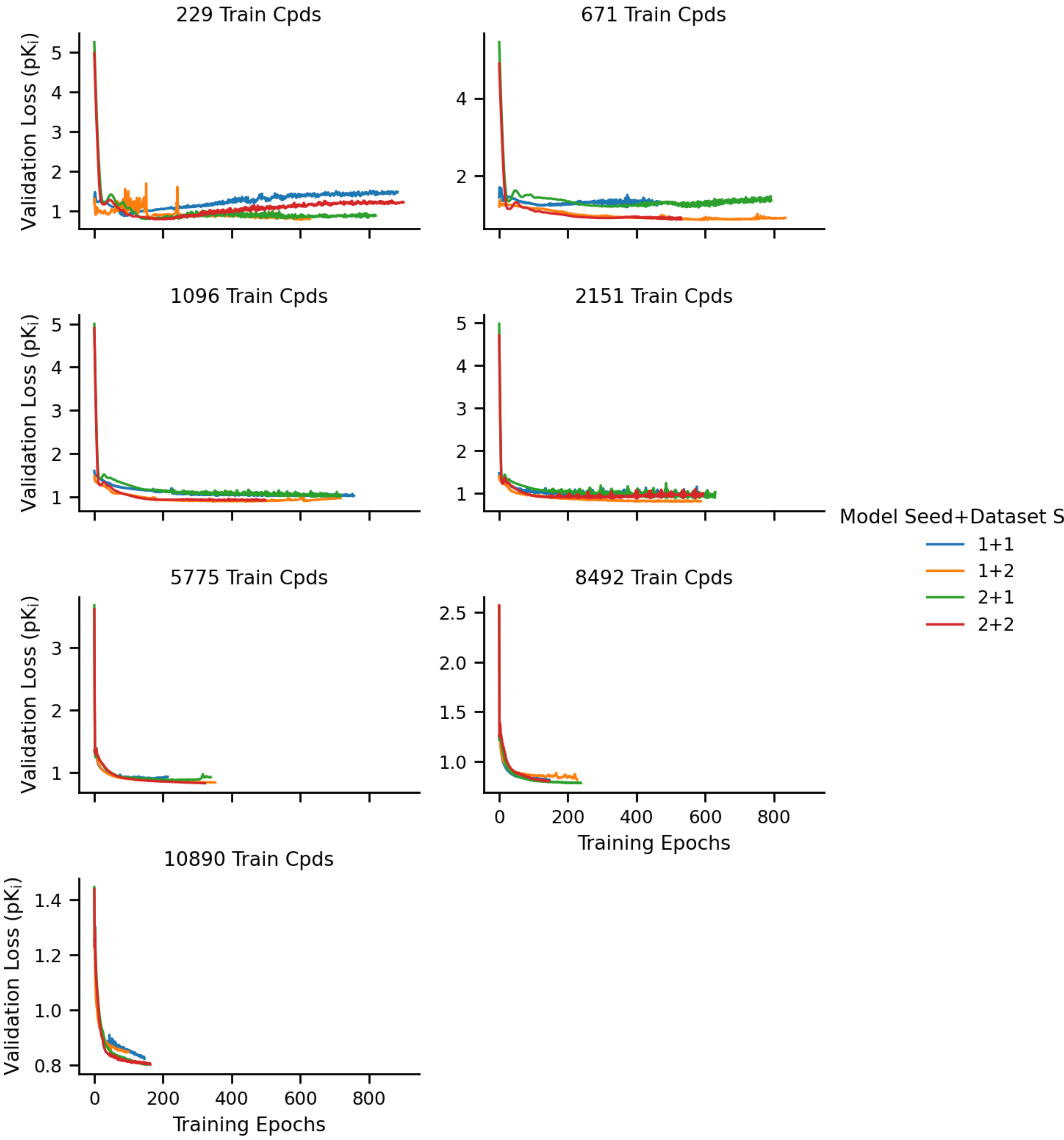


**Figure S11. Training curves for training the 3D E(3)-invariant model on the PDBBind dataset.** Replicates across smaller train set sizes appear to have completed training, as validation loss is not decreasing, while runs with larger train set sizes may still be training.

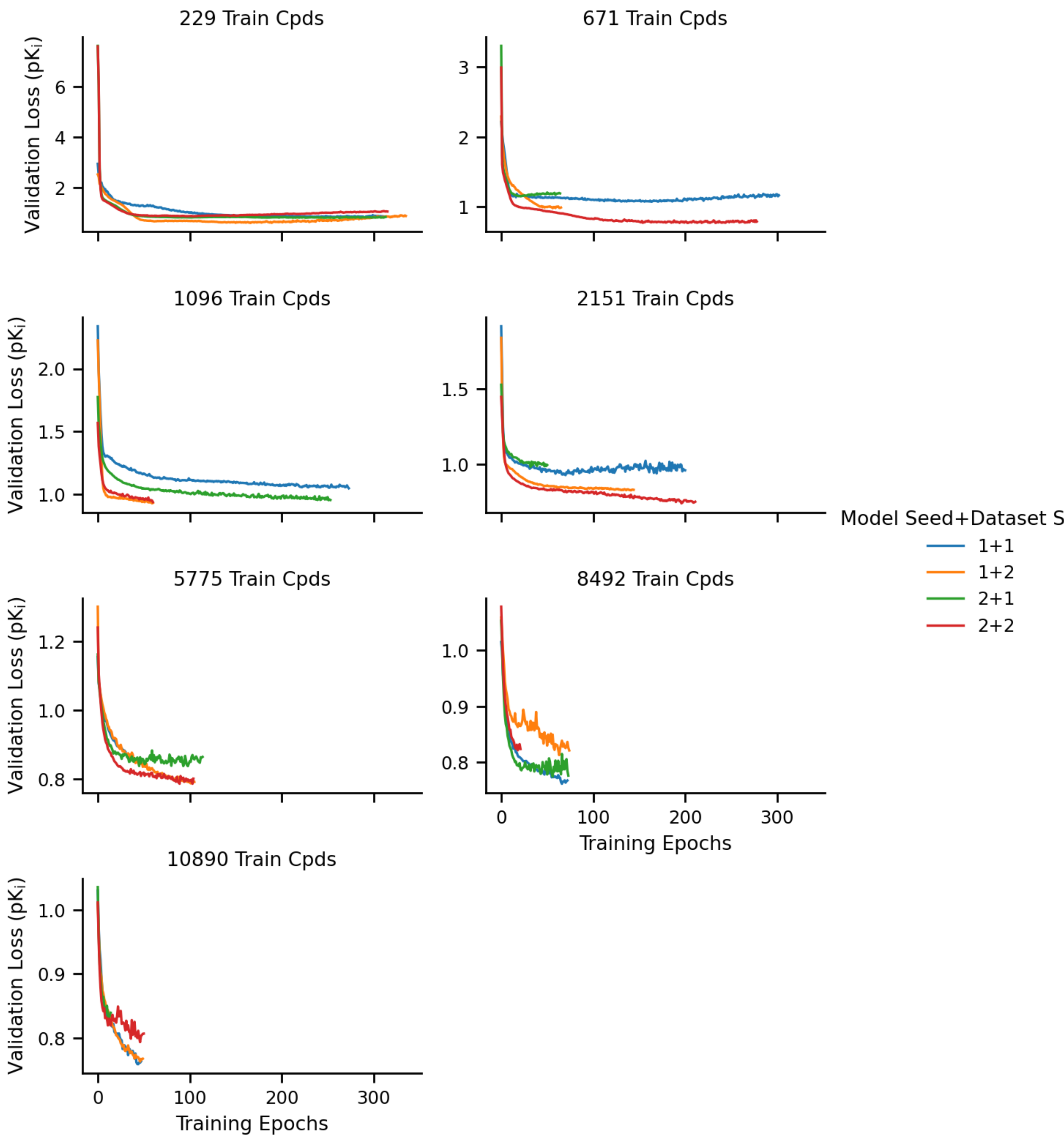


**Figure S12. Training curves for training the 3D E(3)-equivariant model on the PDBBind dataset.** Replicates across smaller train set sizes appear to have completed training, as validation loss is not decreasing, while runs with larger train set sizes may still be training.

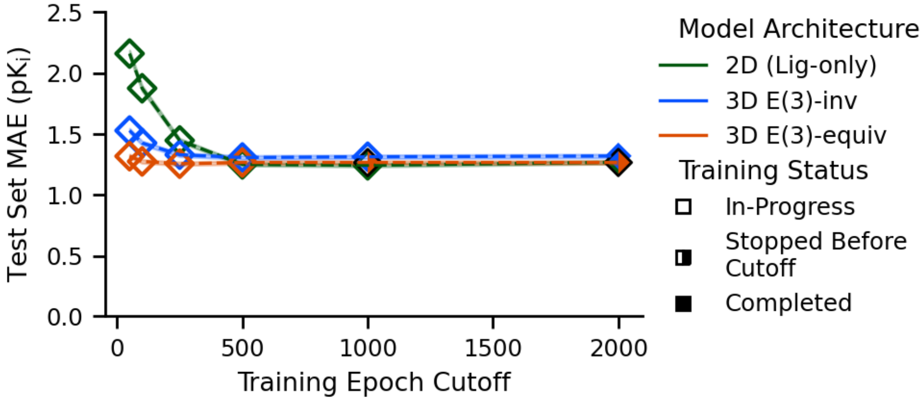


**Figure S13. Model test set performance across different epoch limits of training on a subset of the PDBBind dataset.** Models were trained on 10% (1,136 compounds) of the available PDBBind train set. The smaller train set allows models to train for more epochs in the wall clock cutoff time, but the overall dynamics do not change compared to training on the larger train set for fewer epochs.
